# Cross-Resistance Limits the Ability of Antimicrobial Peptide Combinations to Delay Resistance Evolution

**DOI:** 10.64898/2026.07.05.736553

**Authors:** Bar Maron, Shir Mor, Jonathan Friedman, Zvi Hayouka

**Affiliations:** Institute of Biochemistry, Food Science and Nutrition, The Hebrew University of Jerusalem, Rehovot, Israel; Institute of Environmental Sciences, The Hebrew University of Jerusalem, Rehovot, Israel

## Abstract

**Aims:** Antimicrobial peptide (AMP) combinations have been proposed to delay resistance evolution, but it remains unclear what properties of a peptide pair determine whether a combination reduces resistance evolution relative to its component AMPs used alone. One suggested factor is mode of action, yet this has rarely been tested experimentally. In the current study we have asked whether mode of action or physicochemical similarity between peptides better predicts which combinations delay resistance.

**Methods:** We evolved *Staphylococcus aureus* with six AMPs with reported membrane-targeting and intracellular-targeting activity, individually and in all 15 pairwise combinations. We quantified resistance evolution, cross-resistance and fitness costs across the full AMP panel, and performed whole-genome sequencing on 126 evolved lineages.

**Results:** Resistance varied across AMPs and correlated with peptide chain length, not mode of action. Cross-resistance was associated with physicochemical similarity, and similar peptides selected for overlapping mutations. Most combinations reduced resistance relative to single-AMP treatment, but those whose components shared cross-resistance were less effective, channeling evolution into convergent trajectories that resolve both selective pressures at once. Notably, mode of action did not predict combination outcome.

**Conclusions:** Cross-resistance, not mode of action, is a key factor in determining AMP combination efficacy. Physicochemical distance between peptides may serve as a practical predictor for cross-resistance, enabling selection of AMP combinations that are more likely to constrain resistance evolution.

## Introduction

The emergence of antimicrobial resistance is a significant threat to global public health, causing over a million deaths annually^1,2^. Bacterial pathogens have evolved resistance to virtually all available antibiotics^3,4^, and recent work demonstrates that even next-generation antibiotics face rapid resistance development in vitro^5^. Among the most challenging pathogens is *Staphylococcus aureus* (*S. aureus*) known for its ability to develop resistance to nearly all drug classes while producing toxins, adhesins, and immune-evasion factors that complicate treatment^6–8^.

As current antibiotics become less effective, there is a crucial need for novel therapeutic agents and treatment strategies. Combination therapy is widely used across infectious disease treatment, with the potential to reduce therapeutic doses, decrease side effects, and delay resistance evolution^9–13^.

Antimicrobial peptides (AMPs), natural products of the immune system of most organisms, can offer alternatives to conventional antibiotics^14^. Although diverse in sequence and structure, most AMPs share cationic and amphipathic properties that enable interaction with bacterial membranes^15^. However, accumulating evidence shows that AMPs can also target intracellular components, including DNA, RNA, and the protein synthesis machinery^16^. Notably, AMPs are typically co-produced as mixtures within a single organism rather than deployed individually^17–19^.

Despite their promise as antibiotic alternatives, bacterial resistance to AMPs has been documented across various species and against different peptides^20–22^. The genetic basis of AMP resistance has been characterized in several studies^23,24^, most of which relied on targeted insertion or deletion mutants to identify genes involved in resistance^25,26^. However, these mutants typically confer only modest changes in susceptibility (2 to 4-fold), which may not capture the complexity of resistance that develops under sustained evolutionary pressure. In contrast, experimentally evolved bacteria develop substantially greater resistance through more complex mechanisms involving multiple mutations per strain^27–30^. In previous work, where we evolved *S. aureus* with individual AMPs and their combinations, we identified both common and AMP-specific resistance mutations^31^.

Despite AMPs naturally occurring as mixtures^17–19^, the vast majority of AMP research focuses on individual peptides. This gap extends to resistance evolution, where only a few studies have examined AMP combinations, typically limited to one or two peptide pairs^32–34^. A key open question is what properties of an AMP pair determine whether their combination can delay resistance evolution. One candidate factor is the AMP’s mode of action. Chemical-genetic profiling in *E. coli* revealed that AMPs with different mechanisms of action exhibit more distinct resistance gene fingerprints and more frequent collateral sensitivity than AMPs sharing the same mechanism^35^. Combined with our previous finding that collateral sensitivity between paired AMPs can delay resistance evolution^34^, these results suggest that mode of action could be a critical factor in designing effective AMP combinations. However, an alternative possibility is that the physicochemical properties of AMPs, which shape how bacteria interact with and resist them, may better predict cross-resistance and therefore combination efficacy, regardless of nominal target classification.

To distinguish between these possibilities, we systematically evolved S. aureus with six AMPs representing membrane-targeting and intracellular-targeting mechanisms, both individually and in all 15 pairwise combinations. We addressed three questions:(1) How does resistance evolution vary across AMPs, and can peptide properties predict it? (2) What determines whether an AMP combination succeeds or fails in delaying resistance? (3) What genetic mechanisms underlie cross-resistance between AMPs, and how do these shape combination outcomes? Answering these questions could provide a basis for selecting AMP combinations that delay resistance evolution based on their physicochemical similarity rather than categorical classifications alone.

## Results

### Resistance evolution varies across AMPs and correlates with peptide chain length

To test whether resistance evolves differently to AMPs with different mechanisms of action, we conducted experimental evolution of *S. aureus* with six AMPs: three membrane-targeting (Melittin, Pexiganan, BmKn2) and three intracellular-targeting (Puroindoline, Pleurocidin, Smp24), selected based on antimicrobial activity from a larger panel (**Table S1**). The experimental evolution procedure was based on our previous study, with minor modifications^34^ (see Methods). Briefly, each of six replicate lineages per AMP was exposed to three concentrations (1×, 0.5×, and 0.25× MIC) with selection every four days for the most resistant population (**Fig. 1A**). When the majority of evolutionary replicates (≥4 out of 6) grew at 1× MIC (the highest concentration), the MIC was doubled to maintain selective pressure. The experiment continued for 28 days, constituting approximately 120 generations.

**Figure 1.**
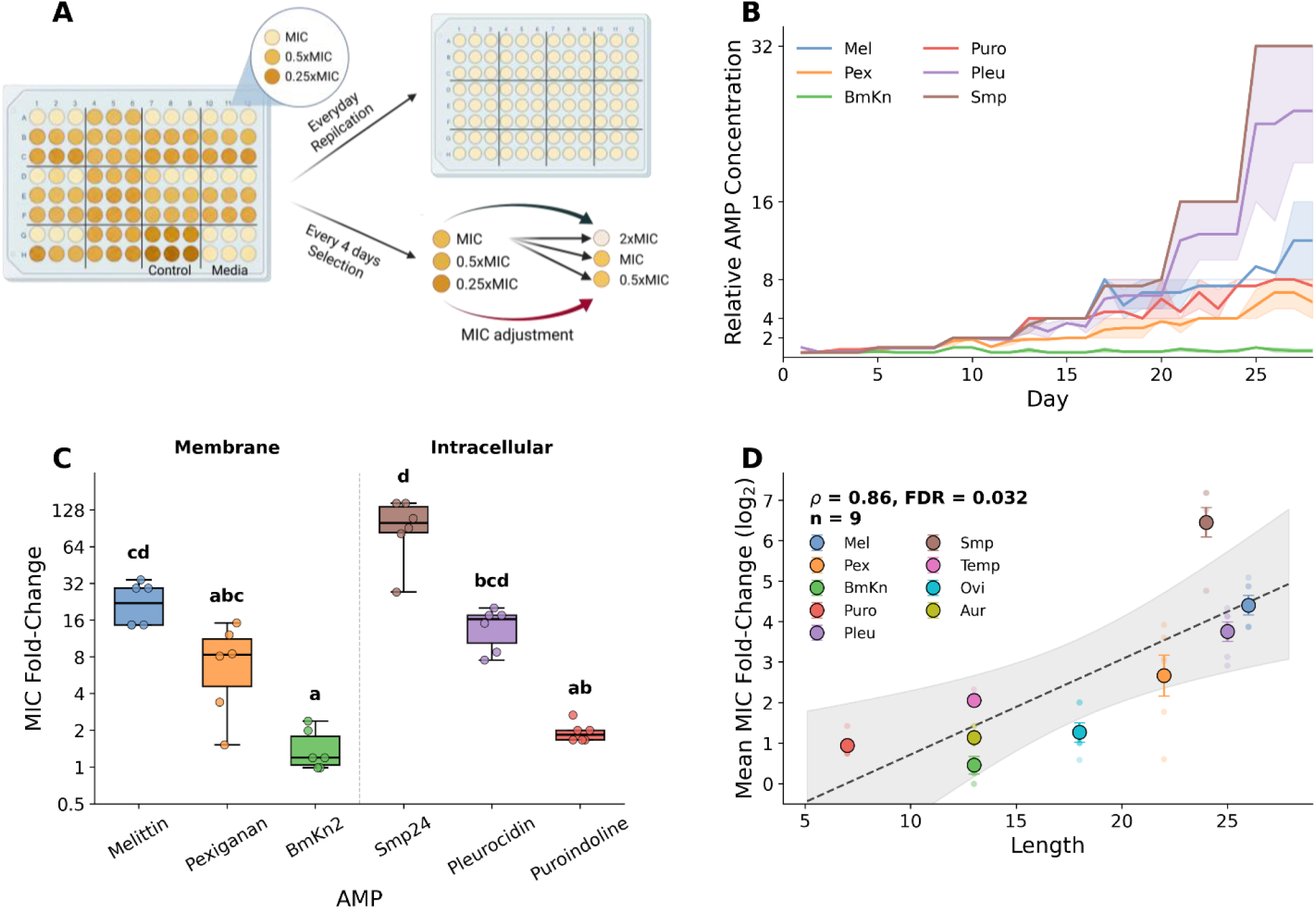
Resistance evolution varies across AMPs and correlates with peptide length. (**A**) Schematic of the experimental evolution protocol. Six evolutionary replicates per AMP treatment were exposed to three concentrations (1×, 0.5×, and 0.25× MIC) in 96-well plates. Cultures were transferred daily into fresh medium, and every four days the most resistant population was selected and propagated across three new concentrations. When ≥4 of 6 replicates grew at 1× MIC, concentrations were doubled. (**B**) Resistance trajectories over 28 days of evolution. Lines show mean relative AMP concentration (normalized to ancestral MIC) across six evolutionary replicates, with shaded areas representing standard error. (**C**) Endpoint MIC fold-change of evolved populations relative to the ancestor, measured under standard conditions. AMPs are grouped by mode of action (membrane-targeting, left of dashed line, and intracellular-targeting, right). Boxes show median and interquartile range, with individual evolutionary replicates as points. Letters indicate post-hoc pairwise comparisons (Dunn’s test with FDR correction, groups sharing a letter do not differ significantly). Overall difference: Kruskal-Wallis H = 56, p = 1.8 × 10⁻⁵. (**D**) Correlation between peptide length and mean resistance (log₂ MIC fold-change) across nine AMPs, including three from a previous study (Temporin-A, Ovispirin, and Aurein 1.2, ref 33). Large circles with error bars show mean ± standard error per AMP. Small faded points show individual evolutionary replicates. Dashed line shows linear regression with 95% confidence interval. Spearman ρ = 0.86, FDR = 0.032.

Resistance remained near baseline for approximately 10 days before increasing at rates that varied substantially among AMPs (**Fig. 1B**). Smp-evolved bacteria grew at concentrations 32-fold higher than the ancestral MIC, whereas BmKn-evolved lines could not grow above the ancestral MIC.

To confirm that resistance was heritable, we isolated populations from the final transfer and determined MIC values under standard conditions (**Fig. 1C**). Resistance levels varied significantly across AMPs (Kruskal-Wallis H = 56, p = 1.8 × 10⁻⁵), ranging from near-ancestral MIC for BmKn to 128-fold increases for some Smp24 replicates. In some cases, endpoint MIC values did not match the concentrations tolerated during evolution. Smp-evolved bacteria grew at 32-fold MIC during evolution but reached up to 128-fold in isolated MIC assays, likely because the evolution protocol capped the maximum tested concentration. Conversely, Puroindoline-evolved bacteria tolerated 8-fold MIC during evolution but showed only approximately 2-fold MIC increases under standard conditions, suggesting that growth at elevated concentrations was partly mediated by phenotypic tolerance rather than stable genetic change.

MIC fold-change did not differ significantly between membrane-targeting and intracellular-targeting AMPs (Mann-Whitney U = 114, n = 18 per group, p = 0.13), indicating that mode of action did not explain the variation in resistance. We therefore asked whether physicochemical properties of the AMPs could. Including data from three additional AMPs characterized in a previous study (Ovispirin, Temporin-A, and Aurein 1.2^34^), we found a strong positive correlation between peptide chain length and mean resistance (Spearman ρ = 0.86, FDR = 0.032, n = 9 AMPs, **Fig. 1D**). Among other properties, hydrophobic moment and aggregation propensity showed nominally significant correlations with resistance (Pearson r = −0.82 and r = 0.75, respectively, raw p < 0.02), but neither survived multiple testing corrections (FDR > 0.05, **Table S2**). Because peptide chain length, aggregation propensity, and hydrophobic moment are partially correlated with each other, we cannot determine from these data alone which property is likely to causally influence resistance evolution.

Resistance evolution was accompanied by fitness costs in the absence of AMPs. Maximal growth rate, growth yield, and area under the growth curve were all negatively correlated with resistance level (Pearson r = −0.67, −0.55, and −0.63, respectively, all p < 0.001, **Fig. S1**), indicating that higher resistance imposed greater costs. Lag time did not correlate with resistance (r = 0.16, p = 0.354). BmKn-evolved lines, which showed minimal resistance, exhibited no significant fitness deficit compared to unevolved control.

### Cross-resistance correlates with physicochemical similarity between AMPs

To characterize the spectrum of cross-resistance, we measured the MIC of all evolved strains against the full panel of six AMPs (**Fig. 2A**). Hierarchical clustering of the resulting profiles revealed two main groups. Melittin and Smp-evolved lines formed a cluster with broad cross-resistance to each other and to BmKn, reaching higher BmKn MICs than lines evolved with BmKn itself. Pleurocidin and Pexiganan-evolved lines formed a second, weaker cluster with mild mutual cross-resistance. BmKn and Puroindoline-evolved lines showed minimal cross-resistance to other AMPs, and several lineages displayed collateral sensitivity to Melittin.

**Figure 2.**
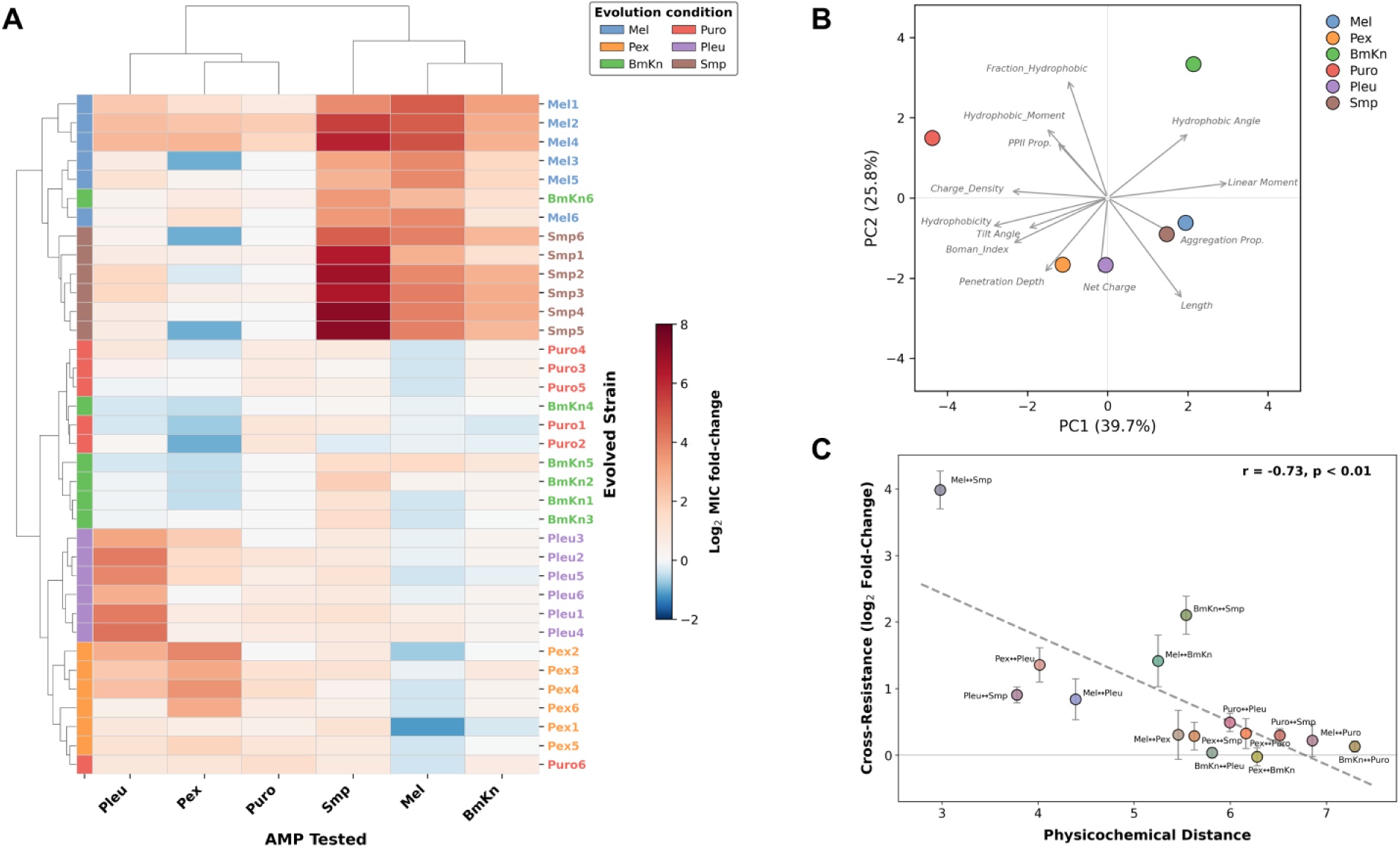
Cross-resistance correlates with physicochemical similarity between AMPs. (**A**) Hierarchical clustermap of cross-resistance profiles. Each row represents an evolved strain tested against the panel of six AMPs (columns). Color intensity indicates log₂ MIC fold-change relative to the ancestor, with red indicating increased resistance and blue indicating collateral sensitivity. Row side colors indicate the AMP used for evolution. Rows and columns are clustered using Euclidean distance with complete linkage. (**B**) PCA biplot of the six study AMPs based on 13 standardized physicochemical descriptors. Points represent individual AMPs, and arrows indicate the direction and magnitude of each descriptor’s contribution to the first two principal components. Pairwise Euclidean distances used in panel C were calculated in the full 13-dimensional descriptor space. (**C**) Correlation between pairwise physicochemical distance and cross-resistance across the 15 AMP pairs. For each pair, cross-resistance was calculated by pooling log₂ MIC fold-change values from both directions and averaging across all biological replicates. Error bars show standard error. Point color represents the RGB average of the two component AMP colors. Dashed line shows linear regression. Pearson r = −0.73, p < 0.01.

These cross-resistance clusters suggested that shared physicochemical properties between AMPs may underlie shared resistance mechanisms. To visualize the physicochemical relationships among the six AMPs, we performed principal component analysis (PCA) on the standardized 13-dimensional property space and plotted the first two components (**Fig. 2B**). Together, PC1 and PC2 explained 65.5% of the total variance (39.7% and 25.8%, respectively). Melittin and Smp clustered in the positive PC1 region, driven by aggregation propensity and peptide length. BmKn shared this positive PC1 range but diverged sharply along PC2, aligning with the hydrophobic angle. Pexiganan and Pleurocidin clustered in the negative PC2 region, associating with net charge and penetration depth. Puroindoline was highly isolated (negative PC1, positive PC2), driven by charge density and hydrophobic moment, and positioned directly opposite the length vector. Notably, these physicochemical groupings align broadly with cross-resistance profiles rather than their respective modes of action. To quantify this relationship, we correlated physicochemical distance with cross-resistance across all 15 symmetrized AMP pairs, where cross-resistance for each pair was calculated by pooling both directions (e.g., Mel→Smp and Smp→Mel). We found a significant negative correlation (Pearson r = −0.73, Mantel test p < 0.01, **Fig. 2C**), indicating that physicochemically similar AMPs tend to share cross-resistance while distant AMPs do not. Leave-one-out sensitivity analysis confirmed the robustness of this correlation (**Fig. S2**).

These cross-resistance patterns raised the question of whether physicochemically similar AMPs also select for overlapping resistance mutations, which would provide a mechanistic basis for the observed correlation.

### Convergent mutations reveal shared and distinct resistance pathways across AMPs

To identify the genetic basis of resistance, we performed whole-genome sequencing on all 36 evolved lineages (six replicates per AMP treatment), sequencing five colonies per endpoint population to capture within-population variation. Three mutations at positions shared with no-peptide control lineages were excluded as likely reference differences or adaptation to growth conditions. After filtering, we identified 155 unique mutations across 33 strains (**Fig. 3A**). Three BmKn-evolved lineages carried no detectable mutations, consistent with the minimal resistance observed in these lines. The mutational spectrum was dominated by missense substitutions (49%), followed by nonsense mutations and frameshift deletions (13% each), with the remainder comprising insertions, in-frame deletions, structural variants, and intergenic changes (**Fig. 3A**). Most mutations (77%) were fixed across all five sequenced colonies, with the remainder reflecting within-population heterogeneity at the experiment endpoint (**Fig. S3, Table S3**).

**Figure 3.**
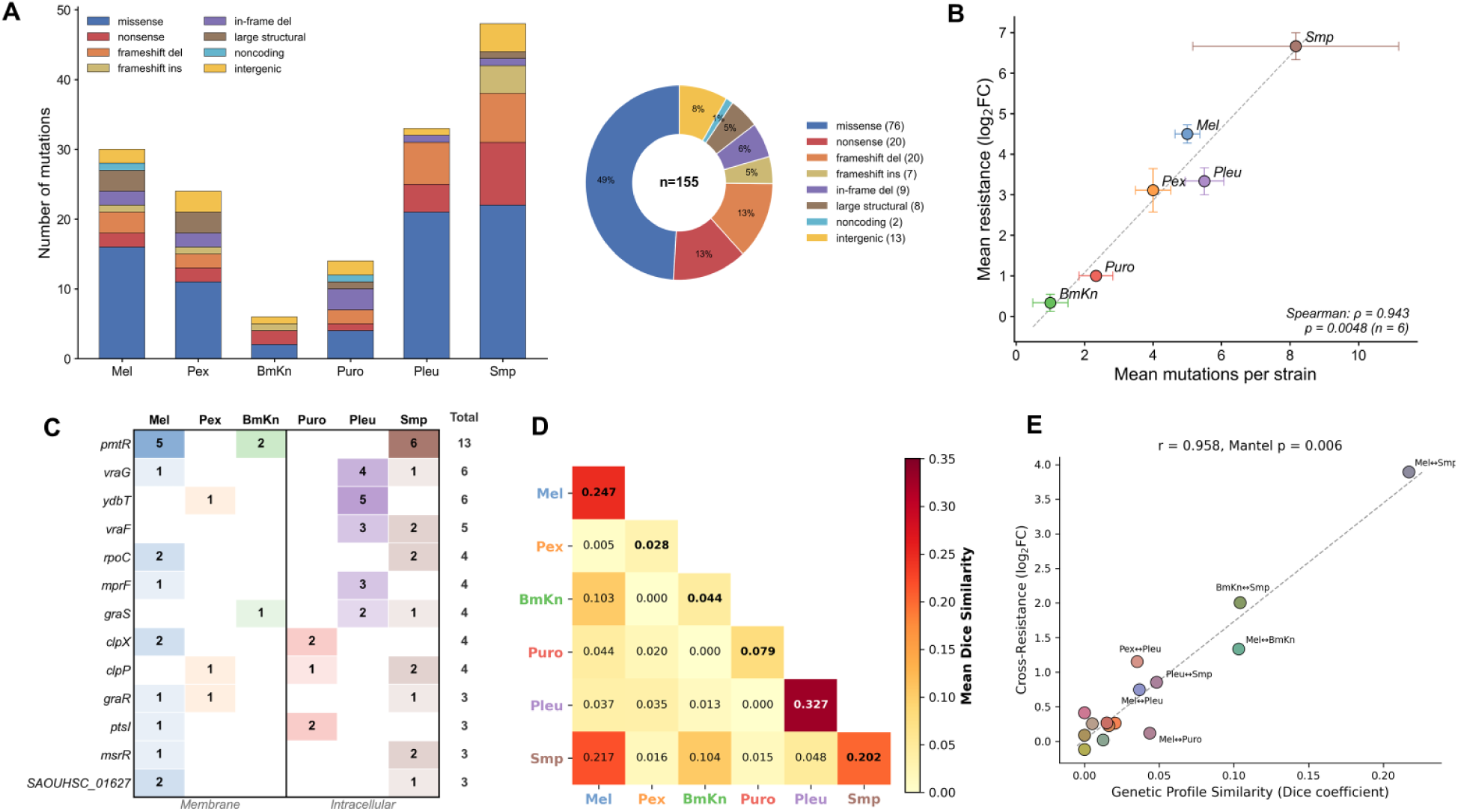
Genetic similarity between AMP-evolved strains correlates with cross-resistance. (**A**) Total number of mutations per AMP treatment (left), colored by variant type. Three BmKn-evolved lineages carried no detectable mutations. Donut chart (right) shows the overall mutational spectrum across all 155 mutations identified in 33 strains. (**B**) Correlation between mean mutation burden per strain and mean resistance level (log₂ MIC fold-change) across the six AMP treatments. Points show treatment means ± standard error. Dashed line shows linear regression. Spearman ρ = 0.943, p = 0.005, n = 6. (**C**) Convergent gene heatmap showing the number of independent lineages carrying mutations in each gene (rows) per AMP treatment (columns), for the genes mutated in three or more lineages. The vertical line separates membrane-targeting (left) and intracellular-targeting (right) treatments. (**D**) Pairwise Dice similarity of mutated gene profiles between and within AMP treatments. Diagonal values represent mean within-treatment similarity. Off-diagonal values represent mean between-treatment similarity. Higher values (darker red) indicate greater overlap in mutated genes. **(E)** Correlation between mean pairwise genetic profile similarity (Dice coefficient) and mean cross-resistance (log₂ fold-change) across all 15 AMP pairs. Each point represents one AMP pair, colored by the RGB average of the two component AMP colors. Labeled points exceed a Dice similarity of 0.03 or a cross-resistance of 0.5. Dashed line shows linear regression. Pearson r = 0.958, Mantel p = 0.006. Significance was assessed by Mantel test (10,000 permutations) to account for non-independence of pairwise values.

Mutation burden varied significantly across treatments (Kruskal-Wallis, p = 2.9 × 10⁻⁴), ranging from a median of 1 mutation per lineage in BmKn-evolved lines to 6 in Smp-evolved lines (**Fig. 3A**). This variation was strongly correlated with resistance level (Spearman’s ρ = 0.943, p = 0.005, **Fig. 3B**), indicating that greater phenotypic resistance required the accumulation of more mutations. One Smp-evolved replicate (Smp2) harbored a *mutS* stop-gain mutation and accumulated 23 mutations, nearly three-fold more than any other strain, without a proportional increase in resistance (log₂FC = 6.76 vs. Smp group mean of 6.45), consistent with a hypermutator phenotype in which most excess mutations are likely neutral passengers. Excluding this strain, the correlation remained significant (ρ = 0.829, p = 0.042), confirming that the relationship is robust and not driven by the outlier.

Thirteen genes were mutated in three or more lineages, together accounting for 41% of all mutations (**Fig. 3C**). The most frequently mutated gene was *pmtR*, encoding a transcriptional repressor of the phenol-soluble modulin transporter (Pmt) operon, with mutations in 13 of 33 strains across three treatments: Smp (6/6), Melittin (5/6), and BmKn (2/3). More than half of these were an identical nonsense mutation (L5*, 7/13), which has also been reported in previous AMP evolution studies^31,32^. The single Melittin-evolved lineage lacking a coding pmtR mutation carried a SNP 41 bp upstream of the gene in its putative regulatory region, suggesting that pmtR inactivation was under strong and consistent selection in Melittin-exposed populations.

The GraXRS/VraFG signaling network was another major target of convergent evolution. This system senses cationic AMPs and coordinates resistance through peptide efflux (VraFG) and membrane charge modification (MprF). Collectively, *vraF*, *vraG*, *graS, graR*, and *mprF* accumulated 22 mutations across five of six treatments, with only Puro-evolved lines showing no mutations in this network. Interestingly, unlike *pmtR* mutations, 86% of these mutations were missense SNPs, suggesting gain-of-function modifications that enhance AMP sensing or efflux rather than loss of protein function. Mutations in *vraF* and *vraG* were particularly enriched in the high-resistance treatments Pleu and Smp.

A second convergent target was *ydbT* (SAOUHSC_02308), encoding a predicted membrane protein whose *B. subtilis* homolog is implicated in envelope stress responses^36^. This gene was mutated in five of six Pleu-evolved lines and one Pex-evolved line, with all six mutations predicted to cause loss of function. The adjacent gene *ydbS* (SAOUHSC_02307) carried additional mutations in two Pex-evolved and one Pleu-evolved lineage (**Table S3**), suggesting that disruption of this locus is specifically selected during adaptation to Pleurocidin and, to a lesser extent, Pexiganan.

The ClpXP protease system was mutated in four strains across three treatments, with Puro-evolved lines contributing three of these (one *clpP*, two *clpX*). All mutations were predicted to cause loss of function, suggesting that reduced ClpXP activity contributes to Puroindoline resistance, potentially through altered regulation of stress response pathways. Finally, *rpoC* (RNA polymerase β’ subunit) was mutated in four strains across Mel and Smp treatments, likely reflecting compensatory adaptation to fitness costs rather than direct resistance.

To quantify overlap between the mutational profiles of evolved lineages, we calculated pairwise Dice similarity of mutated gene profiles within and between treatments (**Fig. 3D, S4**). Within-treatment convergence varied widely, from 0.33 for Pleu to 0.03 for Pex, indicating that some AMPs impose highly reproducible selective pressure while others permit diverse evolutionary solutions. Between-treatment similarity was significantly lower than within-treatment similarity for 9 of 15 comparisons (permutation test, FDR < 0.05). Notably, Melittin and Smp-evolved lines were not genomically distinguishable from each other (Dice = 0.22, FDR = 0.32), driven by shared pmtR inactivation. In contrast, Pleurocidin and Puroindoline shared no mutated genes (Dice = 0) despite both targeting intracellular processes.

To quantify this relationship directly, we correlated mean pairwise genetic similarity between treatments with mean cross-resistance across all 15 AMP pairs. Genetic profile similarity was strongly correlated with cross-resistance (Pearson r = 0.958, Mantel test p = 0.006, **Fig. 3E**), indicating that shared mutations between AMP-evolved lines translate into shared cross-resistance. This correlation was robust to leave-one-AMP-out sensitivity analysis (**Table S4**). Physicochemically similar AMPs also tended to select for overlapping mutations (ρ = −0.616, Mantel p = 0.027 Spearman, **Fig. S2**), though this relationship was weaker and driven primarily by the Mel-Smp and BmKn-Smp pairs.

These patterns of genetic overlap are consistent with the cross-resistance profiles observed in Figure 2: Melittin, Smp, and BmKn share both resistance targets and cross-resistance, while Pleu and Puro do not. Mode of action classification fails to capture these relationships, as the strongest genomic similarity (Mel-Smp) crosses MOA boundaries while the greatest divergence (Pleu-Puro) occurs within the same MOA category. Given that cross-resistance and its genetic basis are shaped by physicochemical similarity rather than mode of action classification, we next asked whether these patterns predict the efficacy of AMP combinations in delaying resistance evolution.

### Cross-resistance rather than mode of action determines AMP combination efficacy

To test whether AMP combinations can delay resistance evolution, we evolved *S. aureus* with all 15 pairwise combinations of the six AMPs. As most combinations showed additive interactions with no synergism or antagonism detected (Table S_), the experiment was performed identically to the single-AMP evolution, with the 1× MIC treatment containing 0.5× MIC of each peptide.

We quantified combination efficacy using two complementary metrics. The relative endpoint resistance (MIC fold-change) of combination-evolved lines to their corresponding single-AMP controls, capturing stable heritable resistance at the experiment endpoint. The relative resistance trajectory compares the area under the daily tolerated concentration curve during evolution to that of the best-performing single AMP, capturing the full resistance trajectory including transient adaptations (**Fig. 4A,B**).

**Figure 4.**
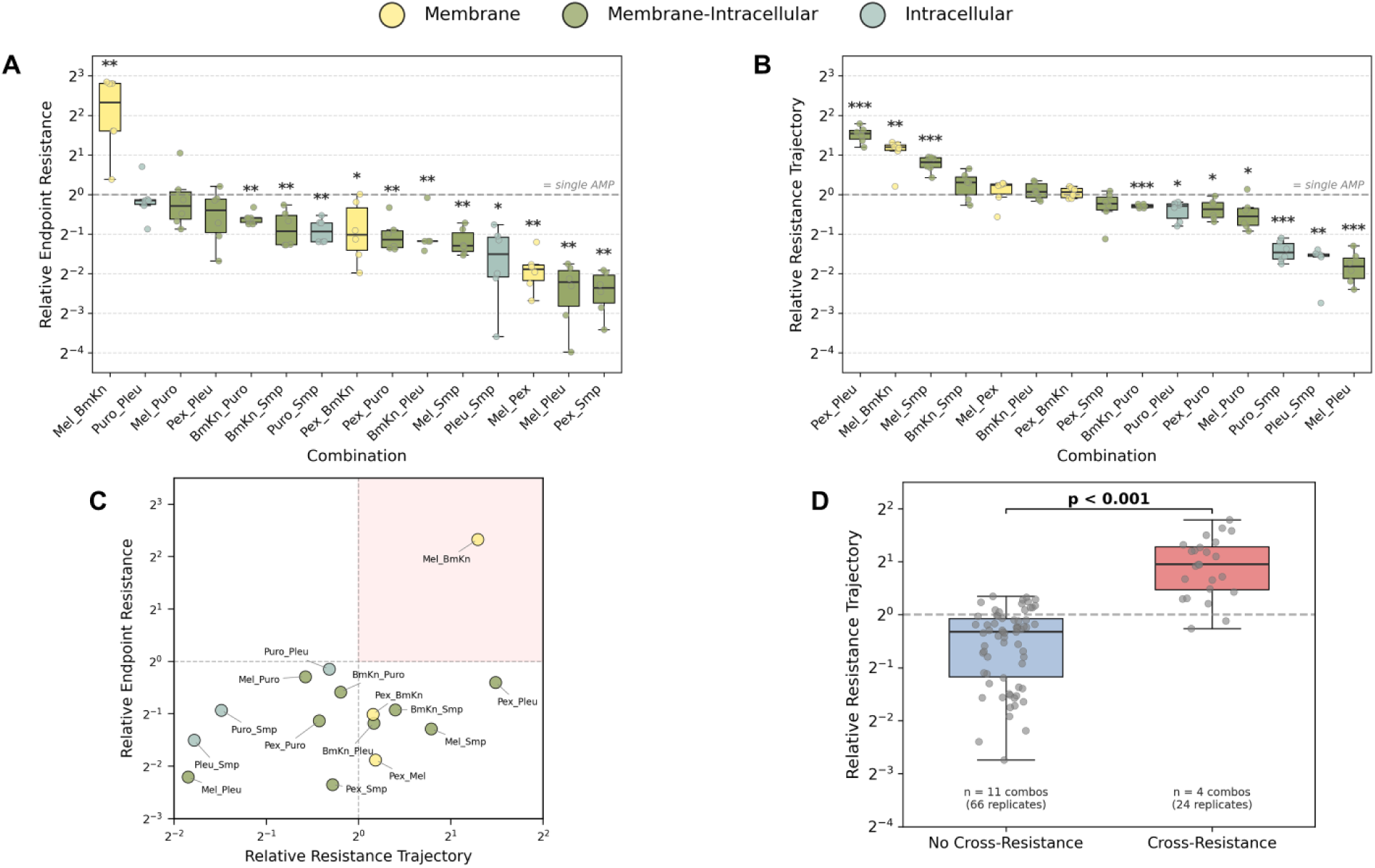
AMP combinations constrain resistance evolution unless component AMPs share cross-resistance. **(A)** Relative Endpoint Resistance for each of the 15 pairwise AMP combinations, defined as the mean ratio of combination-evolved endpoint MIC to the corresponding single-AMP MIC, averaged across the two component AMPs. Values below 1 (dashed line) indicate reduced resistance relative to single-AMP evolution. Combinations are sorted by median Relative Endpoint Resistance. Asterisks denote significant difference from 1 (one-sample t-test on log₂-transformed values, BH-FDR corrected, FDR < 0.05). **(B)** Relative Resistance Trajectory for each combination, defined as the area under the daily tolerated concentration curve of the combination divided by that of the best-performing single AMP. Values below 1 indicate a slower resistance trajectory than the best single AMP. Sorted and annotated as in (A). **(C)** Relationship between the two efficacy metrics. Each point represents one combination (mean across replicates). Dashed lines at 1 on both axes separate combinations that outperformed (below/left) or underperformed (above/right) their single-AMP controls. The red-shaded quadrant indicates combinations that failed by both metrics. Throughout (A-C), colors indicate MOA pairing category: yellow, membrane pair; olive, membrane × intracellular; teal, intracellular pair. **(D)** Relative Resistance Trajectory grouped by cross-resistance status. Combinations were classified as cross-resistant if the mean log₂FC across both testing directions exceeded 1. Each gray point represents one evolutionary replicate. Box plots show median and interquartile range. Mann-Whitney U test performed at the combination level (n = 11 vs. 4 combinations, p < 0.001).

By the relative endpoint resistance, 14 of 15 combinations reduced endpoint resistance relative to single-AMP controls, with 11 reaching statistical significance (**Fig. 4A, S5**). The strongest suppression was observed for Pex-Smp and Mel-Pleu, where combination-evolved lines reached only ∼20% of the single-AMP resistance level. The sole exception was Mel-BmKn, which significantly accelerated resistance. By the relative resistance trajectory, 8 combinations showed lower resistance than the best single AMP, with 7 reaching statistical significance, while 3 performed significantly worse (**Fig. 4B, S6**). Three of the combinations with elevated relative resistance trajectory included BmKn, whose near-baseline single-AMP trajectory left little room for improvement.

The two metrics were correlated but not interchangeable (**Fig. 4C**). For example, Mel-Smp showed a significantly reduced relative endpoint resistance yet an elevated relative resistance trajectory, indicating that bacteria tolerated higher concentrations during evolution but did not retain this resistance as stable MIC increases. Given these complementary perspectives, both metrics are reported throughout.

The only combination that consistently accelerated resistance evolution was Mel-BmKn, with significantly elevated relative endpoint resistance and resistance trajectory (**Fig. 4A,B**). This was driven by increased BmKn resistance: while BmKn-evolved lines alone showed minimal resistance (log₂FC < 0.3), Mel-BmKn-evolved lines reached BmKn MICs more than 8-fold higher than the single-AMP BmKn mean (**Fig. S5**). Because BmKn imposed minimal selective pressure on its own, resistance mechanisms acquired under Melittin exposure likely conferred cross-resistance to BmKn, effectively neutralizing the second component and removing the independent selective pressure needed to constrain resistance evolution. This led us to ask whether cross-resistance between paired AMPs could explain combination outcomes more broadly.

To test this, we classified each combination as cross-resistant if the mean cross-resistance across both directions (A-evolved tested against B and B-evolved tested against A) exceeded log₂FC > 1, corresponding to an average two-fold MIC increase. Four of 15 combinations met this criterion. Combinations with cross-resistance had significantly higher relative resistance trajectory than those without (Mann-Whitney p < 0.001, **Fig. 4D**), indicating that the propensity for cross-resistance between paired AMPs undermines combination efficacy.

In contrast, the mode of action pairing (same or different target) was not associated with combination efficacy by either metric (Kruskal-Wallis, pEndpoint = 0.81, pTrajectory = 0.093; **Fig. S7**), further supporting cross-resistance as the operative predictor of combination outcome rather than nominal target classification.

### Mutational targets under combination therapy are dictated by component AMP identity

To determine whether combination therapy alters resistance mechanisms or recapitulates single-AMP evolutionary trajectories, we sequenced 90 combination-evolved lineages (15 combinations × 6 replicates). We identified 350 mutations affecting 119 genes and 22 intergenic loci. Mutation burden per strain did not differ between combination and single-AMP evolved lineages (median 4.0 vs. 4.0, Mann-Whitney p = 0.74), nor did the distribution of variant types or functional effects (χ² p = 0.30 and 0.26, respectively, **Fig. S8**).

Moreover, the median mutation burden of each combination correlated with the mean mutation burden of its two parent single-AMP treatments (Spearman ρ = 0.74, p = 0.002, **Fig. S8**), indicating that the evolutionary complexity of a combination is largely determined by its component AMPs. At the gene level, the majority of mutations in combination-evolved strains occurred in genes not mutated in either parent single-AMP lineage (median 67% per combination, **Fig. S9**), and no individual gene was significantly enriched or depleted relative to single-AMP therapy after correction (all FDR > 0.05). This novelty may partly reflect incomplete sampling of the single-AMP mutational landscape (6 replicates per treatment), but it indicates that the overall gene-level targets under combination therapy are not simply the union of the two component AMP profiles.

Despite this overall novelty, a small number of key target genes retained strong AMP-specific associations regardless of combination context (**Fig. 5A**). The presence of Melittin or Smp was significantly associated with *pmtR* locus mutations (Fisher’s exact test, 83% and 78% of exposed strains, respectively, FDR < 0.001), extending the pattern observed under single-AMP selection. Conversely, Pleurocidin was significantly depleted for pmtR mutations (19%, FDR = 0.005), and Puroindoline showed a similar trend (25%, FDR = 0.058). Instead, these AMPs were associated with mutations in distinct targets. Pleurocidin was associated with *ydbT* and *ydbS* mutations (39% and 17%, FDR = 0.003 and 0.048 respectively) and Puroindoline with clpX mutations (25%, FDR = 0.017).

**Figure 5.**
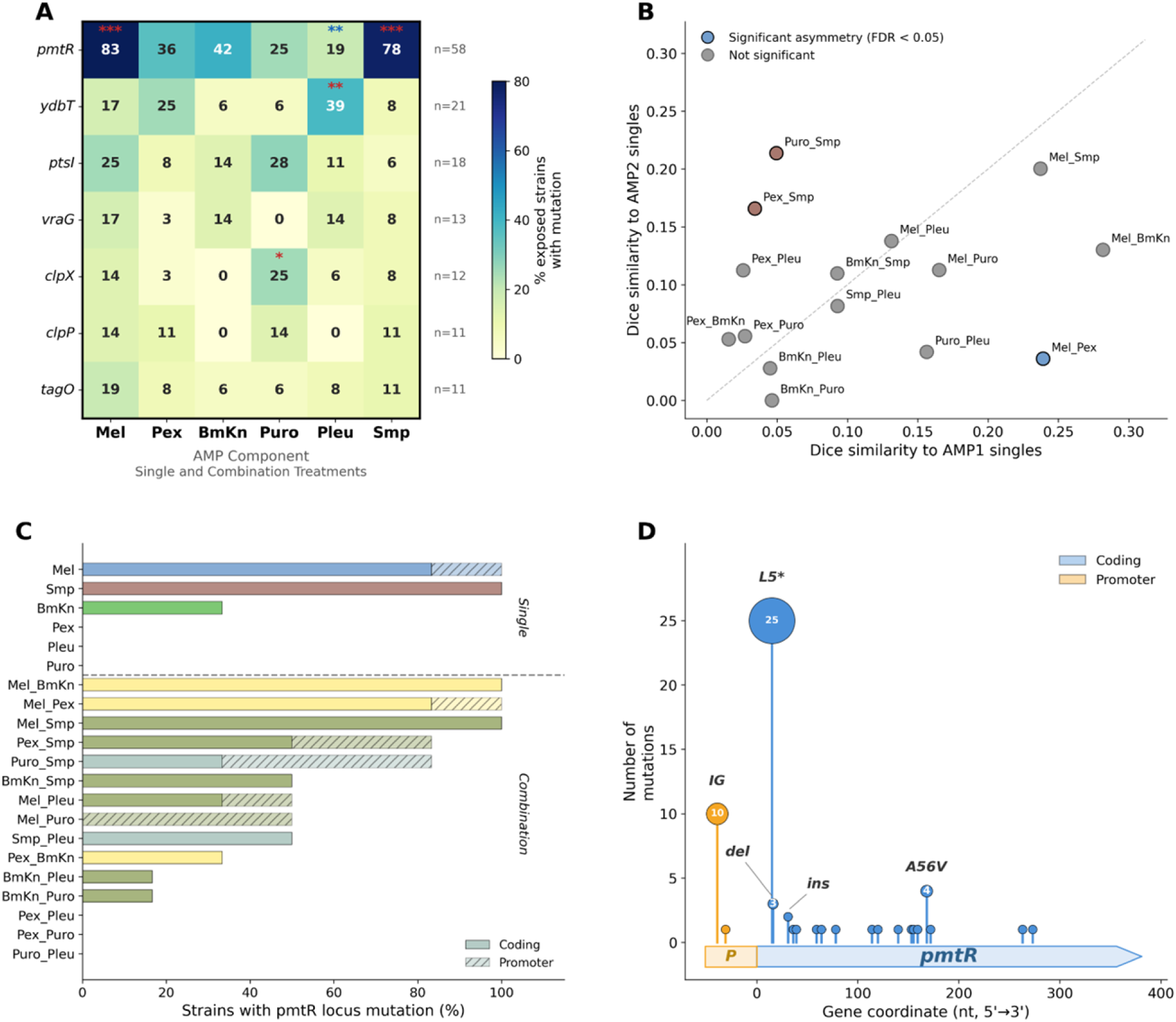
Mutational targets under combination therapy are dictated by component AMP identity. (**A**) Association between individual AMPs and mutations in key resistance genes, pooled across single-AMP and combination-evolved strains. Cell values indicate the percentage of strains exposed to a given AMP (columns) that carry a mutation in each gene (rows). Color scale reflects mutation frequency. Asterisks indicate significant enrichment (red) or depletion (blue) by Fisher’s exact test with Benjamini-Hochberg FDR correction (* FDR < 0.05, ** FDR < 0.01, *** FDR < 0.001). Total number of strains carrying mutations in each gene shown at right (n). **(B)** Genomic similarity of combination-evolved strains to their two parent single-AMP lineages. Each point represents one combination, with x-axis and y-axis showing mean Dice similarity to the first and second parent AMP single-evolved strains, respectively. Points on the diagonal (dashed line) indicate equal similarity to both parents. Colored points indicate combinations with statistically significant parental asymmetry (permutation test, FDR < 0.05), colored by the dominant parent AMP. Gray points indicate no significant asymmetry. (**C**) Prevalence of pmtR locus mutations across all single-AMP (top) and combination (bottom) treatments. Bars show the percentage of strains carrying coding mutations (solid fill) or promoter mutations (hatched fill). Treatments are sorted by prevalence within each category. (**D**) Map of mutations across the pmtR locus. The gene structure is shown at bottom, with the promoter region (P, orange) and coding sequence (blue arrow). Each lollipop represents an independent mutation, with circle size proportional to the number of occurrences. IG denotes intergenic mutations in the putative promoter region.

Integrating mutation data with MIC measurements confirmed that these AMP-specific targets conferred AMP-specific resistance (**Fig. S10**). Strains carrying *pmtR* mutations showed elevated MIC to Melittin, BmKn, and Smp (FDR < 0.001) but not to Pleurocidin or Puroindoline. Similarly, *ydbT* mutations were associated with increased resistance to Pexiganan and Pleurocidin (FDR = 0.005 and 0.012, respectively), and *clpX* mutations with resistance to Puroindoline (FDR < 0.01). Thus, although the overall mutational landscape of each combination is largely unique, the critical bottleneck genes are determined by the identity of the component AMPs and confer resistance specifically to those AMPs.

To quantify the overall genomic relationship between combination and single-AMP evolved strains, we calculated pairwise Dice similarity of mutated gene profiles (**Fig. 5B**). Combination-evolved strains were more similar to their component single-AMP lineages than to unrelated AMPs (mean Dice 0.10 vs. 0.05, Mann-Whitney p = 0.015), confirming that the selective signatures of individual AMPs persist in the combination setting. However, this similarity did not exceed the within-treatment baseline of either parent in any combination (permutation test, all FDR > 0.05), indicating that the observed overlap reflects the predictability of mutational targets under a given selective pressure rather than a fixed set of resistance mutations inherited from one parent. Three of 15 combinations showed significant asymmetry in parental resemblance (permutation test, FDR < 0.05): Mel-Pex strains were more similar to Mel than to Pex (Dice 0.23 vs. 0.04, p = 0.002), while Pex-Smp and Puro-Smp strains more closely resembled Smp (Dice 0.16 and 0.21, respectively). In each case, the dominant parent was the AMP with higher within-treatment convergence among its own replicates, suggesting that AMPs driving more predictable resistance mutations leave a more recognizable signature on combinations.

These patterns were consistent with the combination efficacy results. Mel-BmKn, the only combination that accelerated resistance, showed the highest within-treatment genomic convergence (Dice = 0.64). Melittin’s strong selection for *pmtR* inactivation dominated the combination, driving mutations that simultaneously conferred cross-resistance to BmKn and neutralized BmKn’s contribution. Conversely, combinations pairing AMPs with non-overlapping mutational targets, such as Mel-Pleu (*pmtR* vs. *ydbT*), constrained resistance by requiring mutations in independent pathways. However, non-overlapping targets alone did not guarantee efficacy. Pex-Pleu showed limited benefit, consistent with both AMPs selecting for *ydbT* mutations even in the single-AMP setting. The degree of overlap in AMP-specific mutational targets, rather than mode of action classification, thus determines whether a combination constrains resistance evolution.

The pmtR locus exemplifies this principle as the most frequently mutated target across both datasets, with mutations detected in 58 of 126 total strains (46%; **Fig. 5C**). In the combination dataset, *pmtR* mutations were present in 12 of 15 treatments, absent only in combinations lacking Melittin, Smp, or BmKn (Pex-Pleu, Pex-Puro, Puro-Pleu). Beyond the coding loss-of-function mutations described in the single-AMP analysis, 11 independent mutations targeted the putative promoter region in combination-evolved strains, including a recurrent C→G substitution 41 bp upstream of the start codon observed in 10 independent lineages across 8 treatments (**Fig. 5D**). Together with the L5 nonsense hotspot (25 occurrences), these two positions accounted for 59% of all *pmtR* locus mutations, pointing to extreme convergence on a narrow mutational target. The high prevalence of *pmtR* mutations even in strains with minimal resistance suggests that this locus may function as a low-cost, permissive first step rather than a primary resistance determinant. For example, the lineage BmKn5 carries a *pmtR* mutation but exhibits negligible resistance with a log₂FC < 0.3.

Quantitative analysis of all 126 strains confirmed this interpretation. The two strains carrying *pmtR* as their sole mutation (BmKn5 and BmKn-Puro1) exhibited minimal self-resistance (mean log₂FC = 0.52), statistically indistinguishable from pmtR-negative strains (Kruskal-Wallis p < 0.001 across groups, pmtR-only vs. no-pmtR FDR = 0.53), whereas strains carrying *pmtR* together with additional mutations reached significantly higher resistance (mean log₂FC = 3.00, FDR < 0.001 vs. no-pmtR; **Fig. S11A**). The resistance contribution of *pmtR* was restricted to the Mel-BmKn-Smp physicochemical cluster, with pmtR-positive strains showing significantly elevated MIC against Melittin, BmKn, and Smp (all FDR < 0.001) but not against Pexiganan, Pleurocidin, or Puroindoline (**Fig. S11B**). *pmtR* mutations co-occurred with GraRS/VraFG network mutations more frequently than expected by chance (Fisher’s exact OR = 3.45, p = 0.006), and strains carrying both mutation classes achieved the highest resistance levels (mean log₂FC = 4.32), with each contributing independently (**Fig. S11C**).

### Convergent versus divergent evolution underlies combination failure and success

To examine how cross-resistance shapes combination outcomes at the genomic level, we compared two combinations sharing one component (Melittin) but differing in efficacy.

Mel-BmKn, the only combination that accelerated resistance, showed the highest within-treatment genomic convergence of any treatment in the study (mean Dice = 0.64, **Fig. 6A, S12**). Four of six replicates acquired mutations in the same five genes (*pmtR*, *xdrA*, *vraG*, *ptsI*, and *graS*), and all six carried *pmtR* loss-of-function mutations. This convergence was significantly higher than that of both parent single-AMP treatments (vs. Mel, Dice = 0.25, permutation test FDR = 0.032; vs. BmKn singles, Dice = 0.04, FDR = 0.033), indicating that the combination narrowed rather than diversified evolutionary trajectories. Consistent with this, Mel-BmKn evolved strains reached resistance levels against BmKn (mean log₂FC = 3.3) far exceeding those of BmKn-evolved singles (mean log₂FC = 0.4), demonstrating that Melittin-driven mutations conferred more BmKn resistance than BmKn evolution itself.

**Figure 6.**
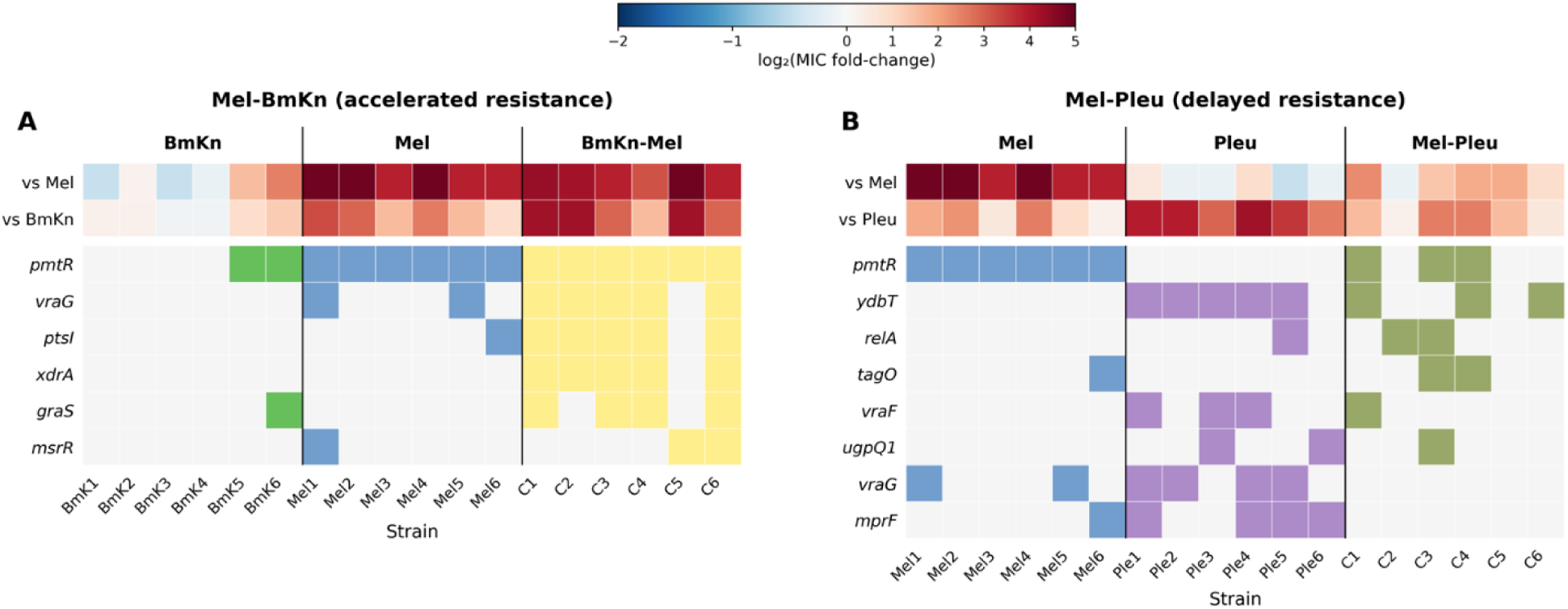
Cross-resistance channels convergent evolution in failing but not successful combinations. (**A**) Mel-BmKn (accelerated resistance) and (**B**) Mel-Pleu (delayed resistance). Upper panels show endpoint MIC fold-change (log₂) of each strain against the two component AMPs, with colors indicating resistance level relative to the ancestral MIC (blue, decreased; red, increased). Lower panels show the presence (colored) or absence (gray) of mutations in genes mutated across the three treatments. Each column represents one evolved lineage from the indicated treatment group. Vertical black lines separate treatment groups. Gene names are ordered by frequency in the combination treatment. Only genes mutated in ≥3 strains are shown (see Fig. S12 for the complete gene set).

Mel-Pleu, one of the most effective combinations, showed the opposite pattern (**Fig. 6B, S12B**). The mean similarity within evolutionary replicates was 0.12, lower than both Mel singles (0.25) and Pleu singles (0.33), and permutation tests confirmed that this combination did not show the elevated convergence observed in Mel-BmKn (FDR > 0.9 for both parent comparisons), reflecting divergent evolutionary trajectories across replicates. The Mel-associated target *pmtR* and the Pleu-associated target *ydbT* were each mutated in three of six replicates but co-occurred in only two, and two replicates carried neither. Endpoint resistance was low (log₂FC = 1.3 vs Mel, 1.5 vs Pleu), approximately 20% of single-AMP levels.

GraXRS/VraFG and *mprF* mutations were absent from the combination despite their prevalence in both parent treatments (*vraG* in 4/6 Pleu and 2/6 Mel singles, *mprF* in 3/6 Pleu and 1/6 Mel singles). Both single-AMP treatments imposed significant fitness costs (**Fig. S1**), and maintaining costly mutations that confer resistance to only one component may be unsustainable under dual selective pressure.

These contrasting cases provide a mechanistic basis for the correlation between cross-resistance and combination efficacy. Overlapping resistance pathways channel evolution into a convergent solution that resolves both selective pressures simultaneously, accelerating resistance. Independent pathways produce divergent trajectories and low resistance, as no single set of mutations can address both AMPs at once.

## Discussion

Recent work has shown that AMP combinations can delay resistance evolution in both *S. aureus* and *P. aeruginosa*, with efficacy depending on factors such as fitness cost, cross-resistance, and mixture diversity^28,34^. However, these studies examined a limited number of defined pairs or used ultra-diverse mixtures, leaving open the question of what properties of an AMP pair determine whether the combination will succeed or fail. By systematically evolving *S. aureus* with all 15 pairwise combinations of six AMPs, we identified a principle linking physicochemical similarity, cross-resistance, and combination efficacy. Broadly, AMP combinations reduced resistance evolution as long as the component peptides did not confer cross-resistance to each other.

One hypothesis for designing effective AMP combinations is that pairing peptides with different modes of action should maximize collateral sensitivity and delay resistance ^35^. This prediction derives from chemical-genetic profiling in *E. coli*, where AMPs with distinct mechanisms showed more divergent resistance gene fingerprints. However, this has not been tested through experimental evolution with actual combination therapy, and the underlying binary classification has weaker foundations than often assumed.

Very few AMPs have sufficient experimental evidence to support a genuinely non-lytic mechanism with substantial intracellular contribution^37^. For most peptides assigned an intracellular mode of action, the classification rests on partial evidence, while membrane-permeabilizing activity has not been carefully excluded. Moreover, the distinction between membrane and intracellular mechanisms is frequently concentration-dependent rather than absolute^16^, and the same peptide can exhibit different mechanisms depending on the target organism and concentration^38^. This is directly relevant to experimental evolution, where bacteria encounter escalating AMP concentrations, meaning the effective mechanism may shift over the course of the experiment. Two of our intracellular-classified peptides illustrate this ambiguity. Pleurocidin derivatives inhibit macromolecular synthesis at sub-lethal concentrations but cause immediate membrane depolarization at higher doses^39^, and Puroindoline-derived peptides bind DNA yet also interact with membranes^40^.

Our genomic data reinforce this conclusion. The strongest between-treatment similarity crossed MOA boundaries (Melittin and Smp24), while the greatest divergence occurred within the same MOA category (Pleurocidin and Puroindoline), and MOA pairing did not associate with combination efficacy. These patterns do not contradict Kintses et al.^35^, whose chemical-genetic approach differs from collapsing mechanism into two categories, and whose findings in Gram-negative *E. coli* may not transfer directly to *S. aureus*. Rather, our results indicate that continuous physicochemical descriptors capture the functional relationships that discrete MOA categories miss, consistent with the argument that physicochemical features enable more accurate classification of AMPs than nominal target-based categories^41^.

The link between physicochemical similarity and combination outcome operates through cross-resistance. Physicochemically similar AMPs selected for overlapping resistance mutations, which in turn produced reciprocal cross-resistance. Combinations pairing AMPs with cross-resistance above a two-fold MIC threshold showed significantly worse efficacy than those without. This finding extends a principle established for conventional antibiotics, where collateral sensitivity between component drugs constrained resistance evolution in *S. aureus* while cross-resistance undermined it, independent of drug interaction type^42^. It is also consistent with our previous work, where Pexiganan-Temporin was an effective combination because Temporin-evolved lines had collateral sensitivity to Pexiganan, whereas Melittin-Temporin failed because the two AMPs conferred cross-resistance to each other^34^. The present study generalizes this pattern across all 15 pairwise combinations and adds a predictive layer absent in prior work. Physicochemical distance between peptides provides a quantitative estimate of cross-resistance likelihood before evolution experiments are conducted. However, cross-resistance functioned as a threshold rather than a gradient in our data. It reliably identified combinations that would fail, but among combinations lacking cross-resistance, the degree of physicochemical distance did not correlate with how effectively the combination delayed resistance. This gap likely reflects additional variables not captured by pairwise cross-resistance alone, including fitness costs of resistance mutations, epistatic interactions between mutations in different pathways, and asymmetries in single-AMP resistance baselines that can distort both efficacy metrics.

The genomic basis of these cross-resistance patterns provides a mechanistic explanation for combination outcomes. Despite the overall novelty of combination mutational landscapes, a small number of bottleneck genes retained strong AMP-specific associations regardless of combination context. *pmtR* was associated with Melittin and Smp exposure, *ydbT* with Pleurocidin, and *clpX* with Puroindoline, and each conferred resistance specifically to its associated AMPs. This pattern is consistent with Jahn et al. framework for conventional antibiotics in *E. coli*, where drug combinations requiring a novel genotype relative to the component drugs showed the lowest evolvability, while combinations whose resistance could be resolved by mutations effective against either component alone showed higher evolvability^43^. In our data, Mel-BmKn exemplifies the latter scenario, as *pmtR* inactivation, driven by Melittin selection, simultaneously conferred cross-resistance to BmKn and channeled all six replicates into a highly convergent evolutionary trajectory. Conversely, in Mel-Pleu, two of six replicates acquired mutations in both *pmtR* and *ydbT*, yet resistance remained low because the GraXRS/VraFG and *mprF* mutations that drove high-level resistance in both parent treatments were absent from the combination. Maintaining costly mutations that confer resistance to only one component may be unsustainable under dual selective pressure, constraining the evolutionary path to high resistance.

The *pmtR* locus was the most frequently mutated target across both datasets, with mutations in 46% of all sequenced strains. PmtR represses the Pmt ABC transporter, which exports phenol-soluble modulins and has been shown to defend *S. aureus* against human AMPs and contribute to virulence during skin infection^44^. Inactivation of *pmtR* likely derepresses Pmt-mediated peptide efflux, consistent with its recurrent selection under AMP pressure in multiple independent studies^31,32,45^. However, our data suggest that *pmtR* inactivation functions as a low-cost permissive first step rather than a primary resistance determinant, as the two strains carrying *pmtR* as their sole mutation exhibited minimal resistance. Higher resistance required additional mutations, particularly in the GraRS/VraFG network, with which *pmtR* co-occurred more frequently than expected by chance. The resistance contribution of *pmtR* was restricted to the Mel-BmKn-Smp physicochemical cluster, conferring no benefit against Pexiganan, Pleurocidin, or Puroindoline. This cluster-specific effect explains why *pmtR* inactivation can simultaneously resolve both selective pressures in cross-resistant combinations like Mel-BmKn, but cannot do so when paired with physicochemically distant AMPs.

Several limitations should be considered. All experiments were conducted in a single *S. aureus* strain under *in vitro* conditions, where the absence of host immune factors, pharmacokinetic fluctuations, and polymicrobial interactions may alter both resistance trajectories and combination outcomes. Whether the physicochemical distance framework generalizes to Gram-negative organisms, whose outer membrane presents a fundamentally different barrier to AMPs, remains to be tested. The framework is built on six AMPs, and although the correlation is robust to leave-one-out analysis, a broader panel would be needed to confirm its generality. BmKn’s near-baseline resistance trajectory creates a floor effect that disproportionately affects both efficacy metrics for the five BmKn-containing combinations, potentially distorting continuous correlations across the dataset. Finally, while the recurrent selection of specific mutations and the existence of strains carrying *pmtR* as their sole mutation provide correlative support for its permissive role, formal genetic validation has not been performed.

This study provides a systematic framework for understanding why some AMP combinations delay resistance evolution while others do not. By testing all 15 pairwise combinations of six AMPs, we show that the majority of combinations effectively reduced resistance relative to single-AMP treatments, consistent with the broader rationale for combination therapy. However, a critical finding is that not all combinations are beneficial. Cross-resistance between component peptides can not only negate the advantage of combination treatment but actively accelerate resistance beyond single-AMP levels, as demonstrated by Mel-BmKn. Physicochemical distance between AMPs provides a quantitative tool to anticipate which pairs are likely to share cross-resistance, enabling rational selection of combination partners before conducting evolution experiments. Importantly, our study examined only pairwise combinations, whereas AMPs naturally occur as complex mixtures within host organisms^17^. How the principles identified here scale to combinations of three or more peptides, and whether increasing mixture complexity further constrains resistance as suggested by random peptide mixture studies^28,34^, remains an important question for future investigation.

## Materials and Methods

### Bacterial strains and media

The ancestor strain used in all experiments was *S. aureus* JLA513 (hla-lacZ hla+, tetracycline resistance, derived from SH1000; kindly provided by Jens Rolff)^46^. Bacterial strains and populations were cultured at 37°C in Mueller-Hinton (MH) (HiMedia) broth or on agar plates. For storage, all bacterial cells in this study were preserved in 25% glycerol at −80°C.

### Peptide synthesis

To select AMPs for experimental evolution, we first screened a panel of candidate peptides with different modes of action for antimicrobial activity against *S. aureus* JLA513 (Table S1). For this initial screen, peptides were synthesized using 9-fluorenylmethoxycarbonyl (Fmoc) solid-phase peptide synthesis (SPPS) on a Liberty Blue synthesizer (CEM, USA). Peptides were cleaved from the resin with a mixture of 95% trifluoroacetic acid (TFA), 2.5% water, and 2.5% triisopropylsilane (TIPS), resuspended in 20% acetonitrile, frozen, and lyophilized. Crude peptides were dissolved in dimethyl sulfoxide (DMSO) and purified by semipreparative reversed-phase high-performance liquid chromatography (RP-HPLC). Peptide mass and purity were verified by matrix-assisted laser desorption ionization time-of-flight mass spectrometry (MALDI-TOF-MS). After initial MIC screening, six AMPs were selected based on confirmed activity: three membrane-targeting (Melittin, Pexiganan, BmKn2) and three intracellular-targeting (Puroindoline, Pleurocidin, Smp24). For the experimental evolution, these six peptides were purchased from Peptide2.0 (USA) at >95% purity, and their activity was confirmed upon arrival and before use in experimental evolution.

### Minimal inhibitory concentration (MIC) determination

Minimum inhibitory concentration (MIC) was determined using a standard broth microdilution protocol^47^. *S. aureus* cells were incubated overnight in MH broth at 37°C with shaking at 200 rpm, diluted 1:100 in fresh MH broth, and grown to an optical density (OD₅₉₅) of 0.1. The culture was then diluted 1:100 to achieve approximately 10⁶ CFU/mL. 100 μL of the adjusted culture was added to each well of a 96-well plate containing two-fold serial dilutions of the tested AMP. Plates were incubated at 37°C for 24 hours and OD₅₉₅ was measured. The MIC was defined as the lowest AMP concentration that inhibited bacterial growth by at least 90%. Each AMP was tested in three technical replicates per plate, and every MIC determination was performed in at least three independent biological repeats. The ancestral MIC for each AMP was calculated as the mean across all biological repeats and used as the reference for fold-change calculations. Resistance of evolved strains was expressed as MIC fold-change (evolved MIC divided by ancestral MIC).

To characterize cross-resistance and collateral sensitivity, each evolved strain was tested against all other five AMPs using the same assay. Ancestral MICs were determined in parallel in each experiment to account for inter-assay variability. Cross-resistance was defined as a mean log₂ fold-change > 1 (corresponding to a two-fold or greater MIC increase) against a non-selecting AMP, and collateral sensitivity as a mean log₂ fold-change < 0.

### Experimental evolution with AMPs

The experimental evolution was performed as previously described, with minor modifications^34^. Prior to evolution with AMPs, a *S. aureus* JLA513 colony was transferred from an MH agar plate into 5 mL MH broth in a 50-mL tube and incubated overnight at 37°C with shaking at 200 rpm. Subsequently, this starter culture was diluted 20-fold into a 1.5-mL Eppendorf tube containing 850 μL MH broth to maintain the same headspace ratio as in the experimental evolution procedure and incubated under identical conditions (37°C, 200 rpm) overnight. These 20-fold dilutions were repeated for seven transfers to habituate the bacteria to the experimental conditions.

Each evolutionary line was exposed to three concentrations of AMP based on its MIC: 1×, 0.5×, and 0.25× MIC (**Fig. 1A**). Experiments were performed in 96-well plates with six parallel lines per AMP treatment. For AMP combination treatments, the effective ratio between AMPs was maintained at 1:1, with the 1× MIC treatment containing 0.5× MIC of each peptide. In each plate, six wells containing bacteria without AMPs served as positive controls, while six wells with medium only served as negative controls to detect contamination. Three lines evolved with rifampicin were included as positive controls for resistance evolution, as *S. aureus* readily develops resistance to this antibiotic.

Daily, 10 μL from the previous plate was transferred into 190 μL of fresh medium containing AMPs. Every four days, bacteria from the highest AMP concentration showing growth were selected and transferred across all three concentrations in a new plate. MIC was doubled when growth (defined as OD₅₉₅ > 0.3) was observed in at least four out of six lines at the 1× MIC concentration. The experimental evolution was conducted for 28 days, representing approximately 120 generations. Before each selection event or MIC increment, samples were collected for glycerol stock preparation (25% glycerol) and stored at −80°C to prevent line extinction. Spot plating was performed on MH agar to confirm bacterial viability before selection.

At the experiment endpoint, each well was spot-plated onto MH agar. For each evolutionary line, bacteria from the highest AMP concentration supporting growth were inoculated into 2 mL MH broth in a culture tube and incubated overnight at 37°C with shaking at 200 rpm. Overnight cultures were transferred to cryovials at a final concentration of 25% glycerol and stored at −80°C. For endpoint MIC determination, a small volume from each glycerol stock was streaked onto MH agar, and approximately three colonies were picked together into a starter culture to capture population-level variation, as some lineages displayed small-colony phenotypes that precluded clean single-colony isolation.

### Fitness cost of evolved bacteria

Growth curves of all single-AMP evolved strains were measured in AMP-free MH broth. Overnight cultures in deep-well 96-well plates (37°C, 800 rpm) were diluted to OD₅₉₅ = 0.1 using a Flowbot automated liquid handling system and transferred to a Synergy microplate reader (BioTek) at 37°C, with OD₅₉₅ readings every 15 minutes for 24 hours. Strains evolved without AMPs were measured in parallel as controls. Each lineage was tested in three independent experiments with two technical replicates. One Pexiganan-evolved lineage (Pex2) was excluded due to insufficient growth. Maximal growth rate and lag time were extracted using Gen5 software (BioTek), growth yield was taken from the final time point, and AUC was calculated by trapezoidal integration over the first 14 hours (Python). All parameters were normalized to the ancestor strain measured in the same experiment.

### Physicochemical properties calculation

Physicochemical properties for all nine AMPs (six from this study and three from a previous study: Temporin-A, Ovispirin, and Aurein 1.2^34^) were obtained from the Database of Antimicrobial Activity and Structure of Peptides (DBAASP)^48^. Four additional properties were calculated from the amino acid sequences using the modlAMP Python library^49^: charge density, hydrophobic moment (Eisenberg consensus scale, α-helical geometry at 100° per residue), fraction of hydrophobic residues, and Boman index. Peptide length was calculated directly from the amino acid sequence. Three DBAASP-derived properties were excluded due to redundancy: isoelectric point (near-zero variance), amphiphilicity index (Spearman ρ = 1 with normalized hydrophobicity), and disordered conformation propensity (ρ = −1 with normalized hydrophobicity). The final feature set comprised 13 physicochemical descriptors (8 from DBAASP, 4 from modlAMP, and peptide length). For correlation analyses between individual properties and resistance, features with coefficient of variation below 0.10 across the tested AMP set were excluded prior to testing.

### Pairwise physicochemical distance

Pairwise physicochemical distances between the six study AMPs were calculated as Euclidean distances in the standardized 13-dimensional feature space. All descriptors were standardized to z-scores (mean = 0, SD = 1) across the six study AMPs using StandardScaler (scikit-learn). Pairwise Euclidean distances were computed from the standardized feature vectors using scipy.spatial.distance.pdist (SciPy). Hierarchical clustering of the distance matrix was performed using complete linkage for visualization (**Fig. 2B**).

### Combination efficacy calculation

Combination efficacy was quantified using two complementary metrics. The relative endpoint resistance, adapted from the Evolvability Index^50^, compares endpoint resistance of combination-evolved lines to their corresponding single-AMP controls. For a combination of AMPs A and B, the relative endpoint resistance for each combination-evolved replicate is:

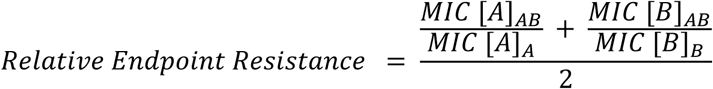

where *MIC* [*A*]*_AB_* is the MIC of the combination-evolved strain against AMP A, *MIC* [*A*]*_A_* is the mean MIC of single-AMP A-evolved strains against AMP A, and equivalently for AMP B. All MIC values are expressed as fold-change relative to the ancestor. A relative endpoint resistance below 1 indicates that the combination reduced endpoint resistance relative to single-AMP evolution.

The relative resistance trajectory compares the resistance trajectory during evolution of each combination replicate to that of the best-performing single-AMP component. For each combination replicate, the area under the daily tolerated concentration curve (normalized to ancestral MIC) was calculated by trapezoidal integration and divided by the mean AUC of whichever single-AMP component showed the least resistance evolution (lowest mean AUC). A relative resistance trajectory below 1 indicates that the combination accumulated resistance more slowly than the best single AMP.

For both metrics, each combination replicate yielded an independent value (n = 6 per combination). Significance was assessed using one-sample t-tests on log₂-transformed values (*H*_₀_: mean = 0, equivalent to testing whether the untransformed ratio differs from 1), with Benjamini-Hochberg FDR correction across the 15 combinations. Log₂ transformation was used because it symmetrizes the ratio scale (values of 2 and 0.5 become equidistant from 0) and improves normality of the test statistic.

### Pairwise similarity analysis

Genomic similarity between evolved strains was quantified using the Dice coefficient on binary gene presence/absence profiles. For each strain, mutated genes were encoded as a binary vector (1 = at least one mutation, 0 = none). Intergenic mutations located within 50 bp of a gene were assigned to that gene’s locus, while more distant intergenic mutations were treated as independent loci. For multi-gene deletions, each affected gene was counted individually. Strains with no detected mutations (three BmKn-evolved lineages) were included as all-zero vectors, yielding Dice = 0 for all their comparisons. Unless stated otherwise, all permutation tests used 10,000 permutations and p-values were corrected using Benjamini-Hochberg FDR. Complete test results are provided in Table S4.

Within-treatment convergence was quantified as the mean pairwise Dice coefficient among replicates of the same treatment. To test whether genetic signatures were distinguishable between single-AMP treatments, permutation tests compared the difference between mean within-treatment and mean between-treatment Dice similarity for all 15 treatment pairs. To assess whether combination-evolved strains retained parent AMP genomic signatures, Dice similarity was calculated between each combination strain and each single-AMP strain, averaged separately for parent and non-parent comparisons, and the difference was tested using a Mann-Whitney U test across the 15 combinations. To test whether combination-to-parent similarity exceeded the baseline within-parent similarity, the mean combination-to-parent Dice coefficient was compared against the mean within-treatment Dice of the corresponding single-AMP group using permutation tests (30 tests: 15 combinations x 2 parents). Parent dominance asymmetry was assessed by comparing the difference in mean Dice similarity to each parent using two-sided permutation tests (15 tests). Finally, to test whether combination treatment produced greater evolutionary convergence than single-AMP treatment, within-treatment Dice similarity of each combination was compared to that of each parent AMP using one-sided permutation tests (30 tests: 15 combinations x 2 parents).

### Whole-genome sequencing

To identify mutations underlying resistance, we performed whole-genome sequencing on all evolved lineages (36 single-AMP and 90 combination-evolved). For each lineage, five colonies were picked from streak plates of the endpoint glycerol stocks, pooled, and grown overnight in MH broth. Genomic DNA was extracted using the NORGEN Bacterial Genomic DNA Isolation Kit, with the addition of 4 μL lysostaphin (1 mg/mL) to enhance cell lysis. Sequencing libraries were prepared by Macrogen (Singapore) using the TruSeq Nano DNA Library Prep Kit (350 bp insert size) and sequenced on an Illumina NovaSeq X Plus platform, generating 151 bp paired-end reads with a target output of 2 Gb per sample, yielding approximately 1,300× average genome coverage.

Variants were called using breseq (v0.39.0) in polymorphism mode against the *S. aureus* JLA513 reference genome (2,720,945 bp). Breseq was run with default parameters and a polymorphism frequency cutoff of 5%. In post-processing, mutations were further filtered to retain only variants present at ≥20% frequency across the five pooled colonies, and variants flagged for strand bias were removed. Three mutations at positions shared with no-peptide control lineages were excluded: one in *agrC* that was present across all samples, consistent with a reference genome difference, and two at positions recurring in multiple control lineages, likely reflecting adaptation to growth conditions rather than AMP resistance. Gene annotations were derived from the JLA513 reference genome with product names assigned based on UniProt common nomenclature.

### Statistical analysis

All statistical analyses were performed in Python (v3.12.7) using SciPy (v1.13.1) and statsmodels (v0.14.2). Non-parametric tests (Kruskal-Wallis, Mann-Whitney U) were used for comparisons involving non-normally distributed data. One-sample t-tests were used to test whether log2-transformed efficacy ratios differed significantly from zero. Chi-square tests of independence were used to compare categorical distributions (variant types, functional effects) between datasets. Correlations were assessed using Spearman rank correlation for ordinal relationships and Pearson correlation for linear relationships. Fisher’s exact tests (two-sided) were used to test associations between AMP exposure and gene mutations, constructed as 2×2 contingency tables comparing mutation frequency in AMP-exposed versus unexposed strains. Tests were restricted to genes mutated in at least six independent strains, corresponding to the number of biological replicates per treatment. Mantel tests were used to assess correlations between pairwise distance or similarity matrices while accounting for the non-independence of values derived from the same set of AMPs. For each test, the observed correlation (Pearson or Spearman, as specified) between upper-triangle elements of two 6×6 matrices was compared to a null distribution generated by simultaneously permuting rows and columns of one matrix (10,000 permutations). Two-tailed p-values were calculated as twice the minimum of the proportion of permuted values exceeding or falling below the observed correlation. Leave-one-out sensitivity was assessed by excluding each of the 15 AMP pairs in turn and recalculating the Pearson correlation (Table S4). Multiple testing correction was applied using the Benjamini-Hochberg method throughout, with a false discovery rate threshold of 0.05. Complete statistical test results are provided in Table S4. Specific tests and sample sizes are described in the relevant Methods subsections and figure legends.

