## Supporting information (Figures S1-12) for "Cross-Resistance Limits the Ability of Antimicrobial Peptide Combinations to Delay Resistance Evolution"

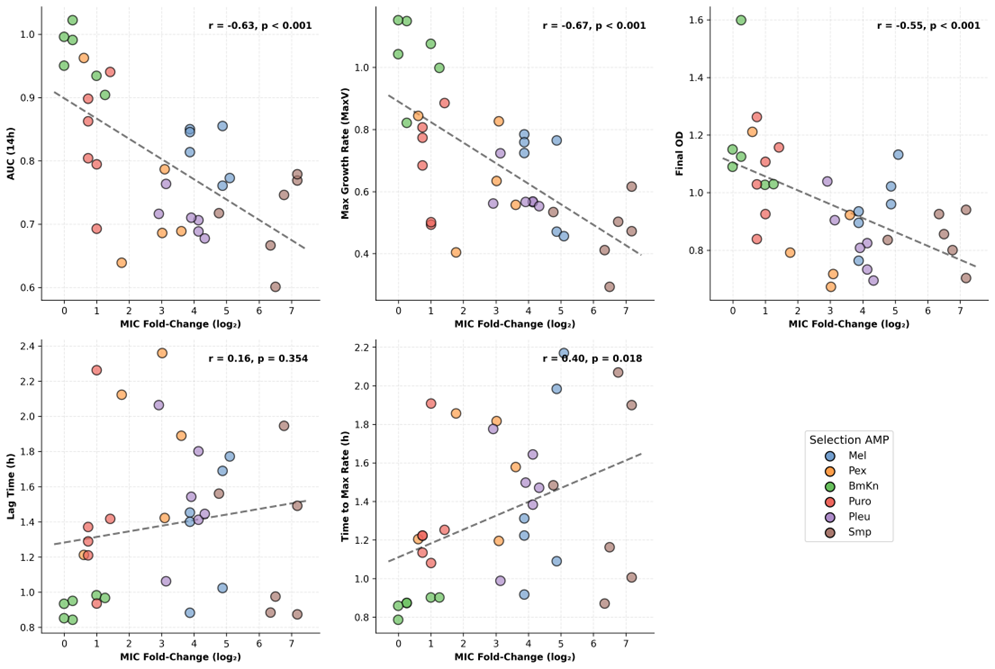
**Supporting information**

**Figure S1. Fitness costs correlate with resistance level.** Correlation between MIC fold-change (log₂) and five growth parameters measured in AMP-free medium: area under the growth curve at 14 hours (AUC 14h), maximal growth rate (MaxV), final optical density (Final OD), lag time, and time to maximal growth rate. Each point represents one evolutionary replicate, colored by selection AMP. Dashed lines show linear regression. Pearson r and p-values are shown for each comparison. Growth rate, AUC, and final OD were negatively correlated with resistance (all p < 0.001), while lag time showed no significant correlation (r = 0.16, p = 0.354).

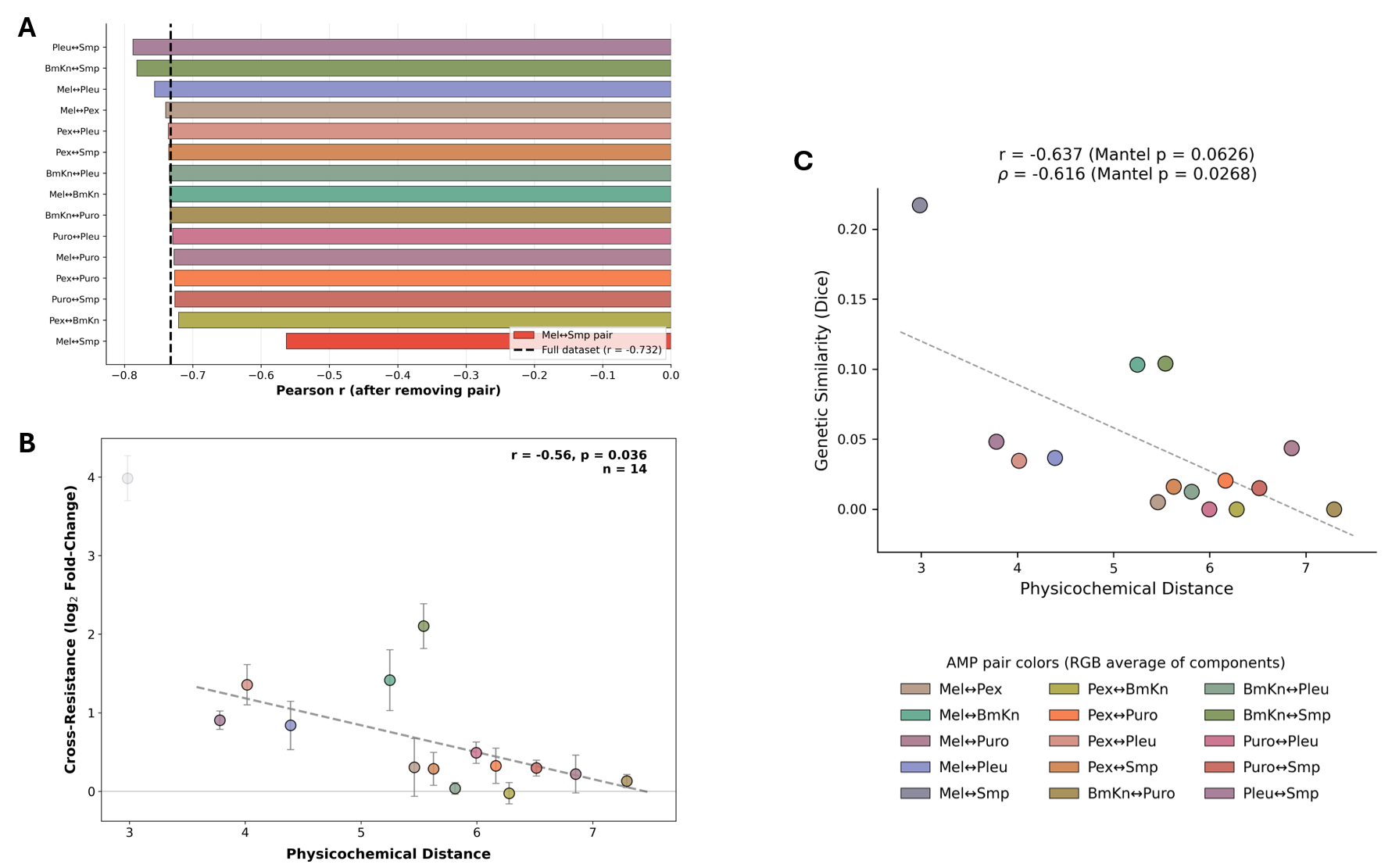

**Figure S2. Leave-one-out sensitivity analysis for the physicochemical distance and cross-resistance correlation.** (**A**) Leave-one-out sensitivity analysis for the physicochemical distance vs. cross-resistance correlation. Each bar shows the Pearson correlation coefficient after removing one symmetrized AMP pair from the full dataset (r = −0.73, dashed line). The Mel↔Smp pair (red) has the largest influence on the correlation. All leave-one-out correlations remain negative. (**B**) Physicochemical distance vs. cross-resistance after removing the Mel↔Smp pair (n = 14). The correlation remains significant (Pearson r = −0.56, p = 0.036), confirming that the relationship is not driven by a single AMP pair. (**C**) Physicochemical distance vs. genetic similarity (Dice coefficient) across the 15 AMP pairs. The correlation is significant by Spearman rank correlation (rho = −0.62, Mantel p = 0.027) but not by Pearson (r = −0.64, Mantel p = 0.063), reflecting the influence of the Mel↔Smp outlier on the linear fit. In all panels, each point represents one symmetrized AMP pair with cross-resistance calculated by pooling both directions, and point colors represent the RGB average of the two component AMP colors (see legend). Error bars in (A) and (B) show standard error.

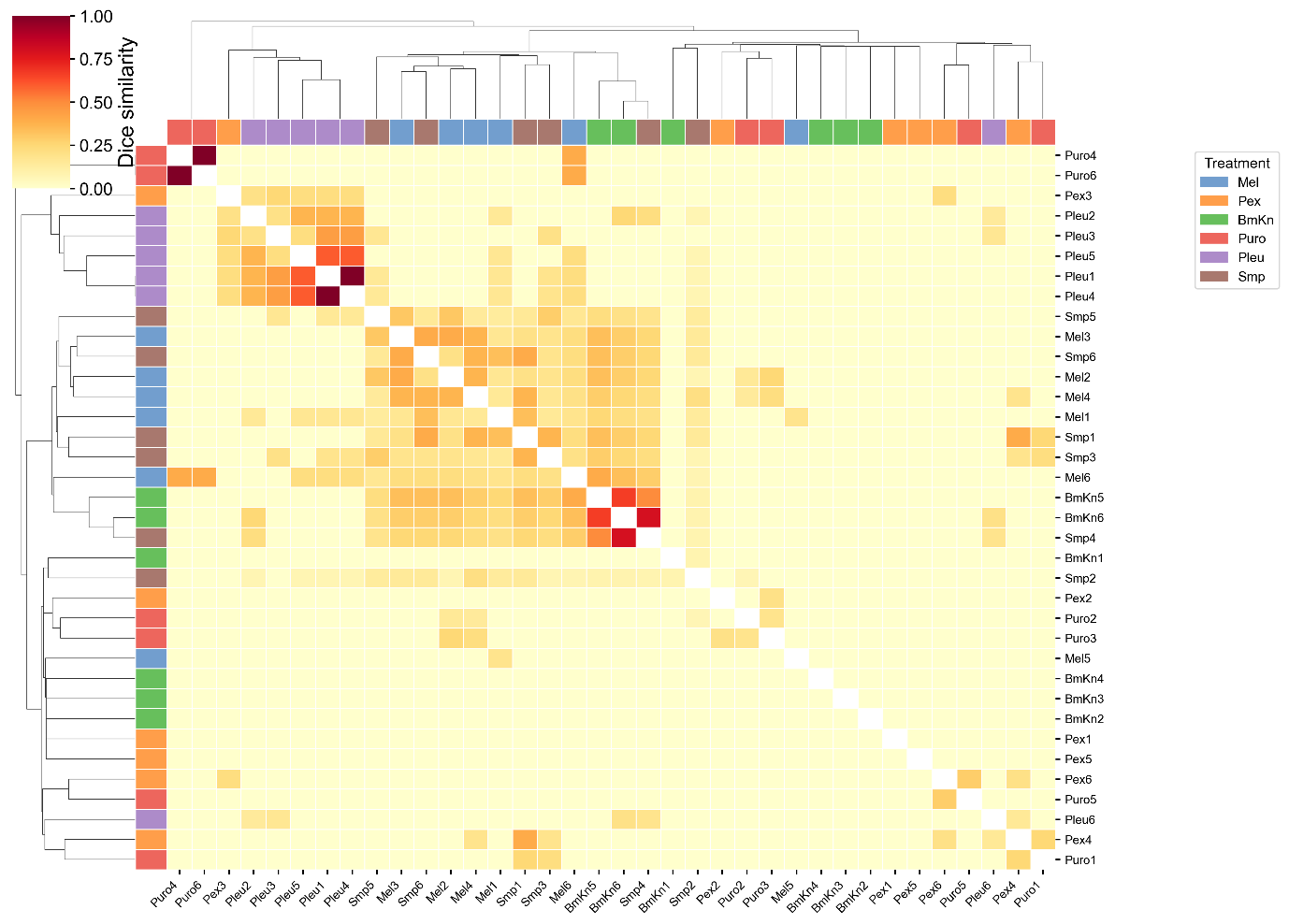

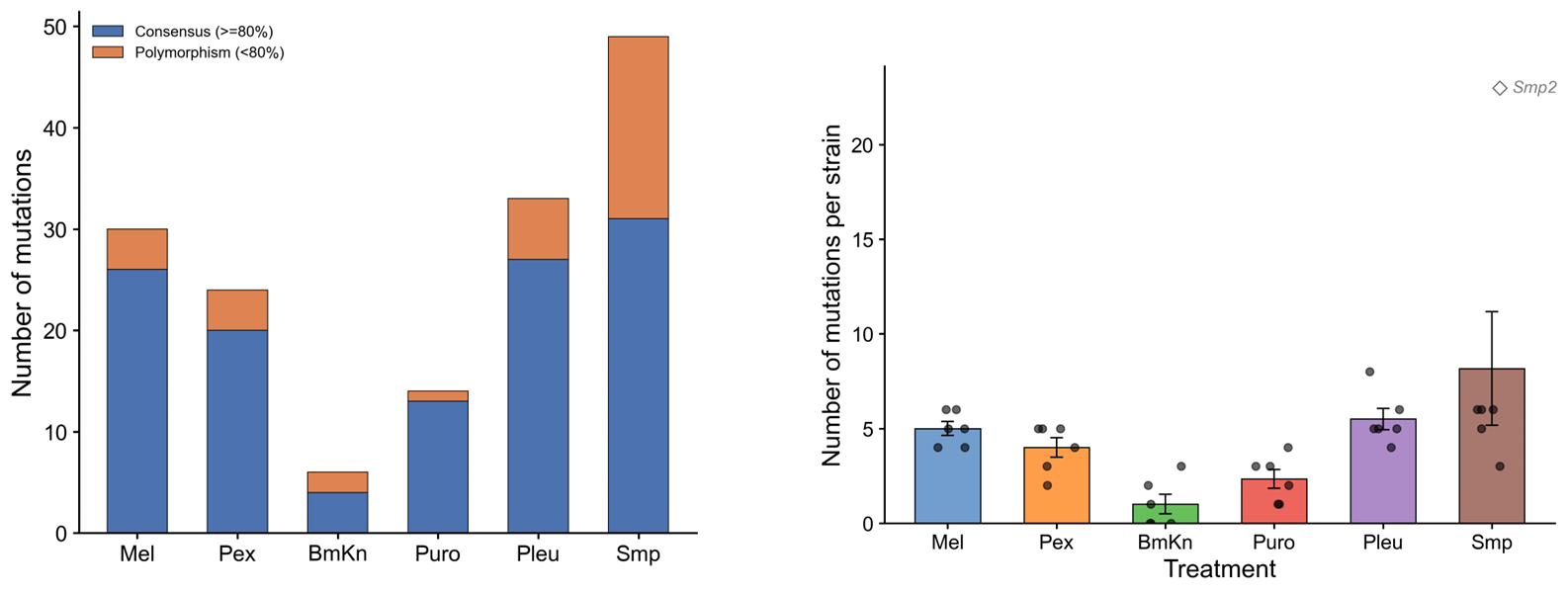

**Figure S3. Mutation fixation and per-strain mutation burden in single-AMP evolved lineages.** (**Left**) Total number of mutations per treatment, classified as consensus (≥80% frequency across sequenced colonies, blue) or polymorphism (<80%, orange). Most mutations were fixed, with Smp showing the highest proportion of polymorphic mutations. (**Right**) Number of mutations per strain across treatments. Bars show means with standard error, and individual strains are shown as points. The Smp2 hypermutator (open diamond, 23 mutations, *mutS* stop-gain) is marked separately.

**Figure S4. Strain-level Dice similarity of mutated gene profiles across single-AMP evolved lineages.** Pairwise Dice similarity between all 33 evolved strains based on shared mutated genes. Diagonal cells (self-comparisons, Dice = 1.0) are masked. Row and column side colors indicate the AMP used for evolution. Hierarchical clustering reveals that strains evolved under the same AMP tend to cluster together, particularly for Pleu and Mel/Smp, while Pex and Puro strains show more dispersed profiles consistent with their lower within-treatment convergence (Fig. 4D)

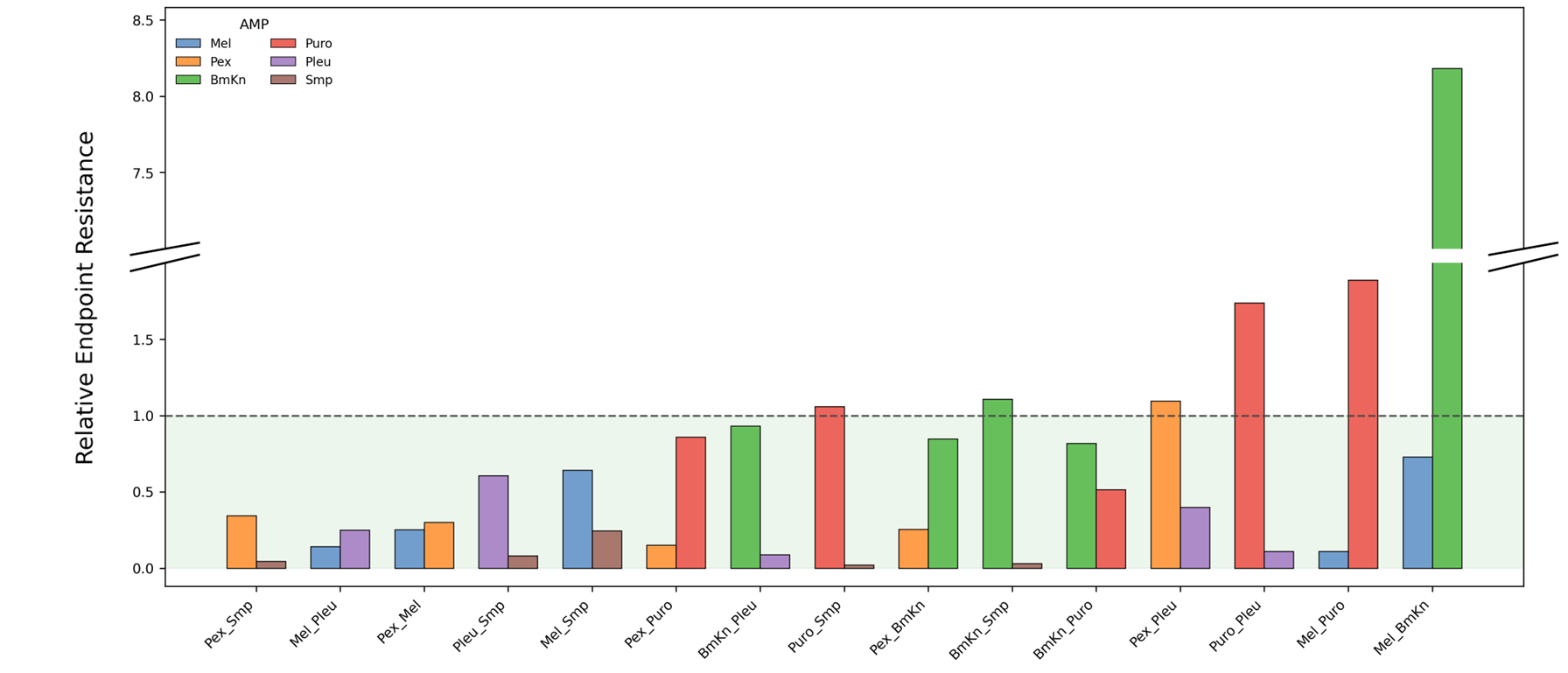

**Figure S5. Per-AMP relative endpoint resistance for each combination.** Each combination is represented by two bars, one for each component AMP, showing the ratio of combination-evolved MIC to the corresponding single-AMP MIC. Bar colors indicate the AMP being evaluated. Values below 1 (dashed line) indicate that the combination reduced resistance to that AMP relative to single-AMP evolution. The green-shaded region highlights values below 1. Combinations are sorted by overall Evolvability Index. Asymmetric bar heights within a combination indicate that one component AMP's resistance was suppressed more than the other.

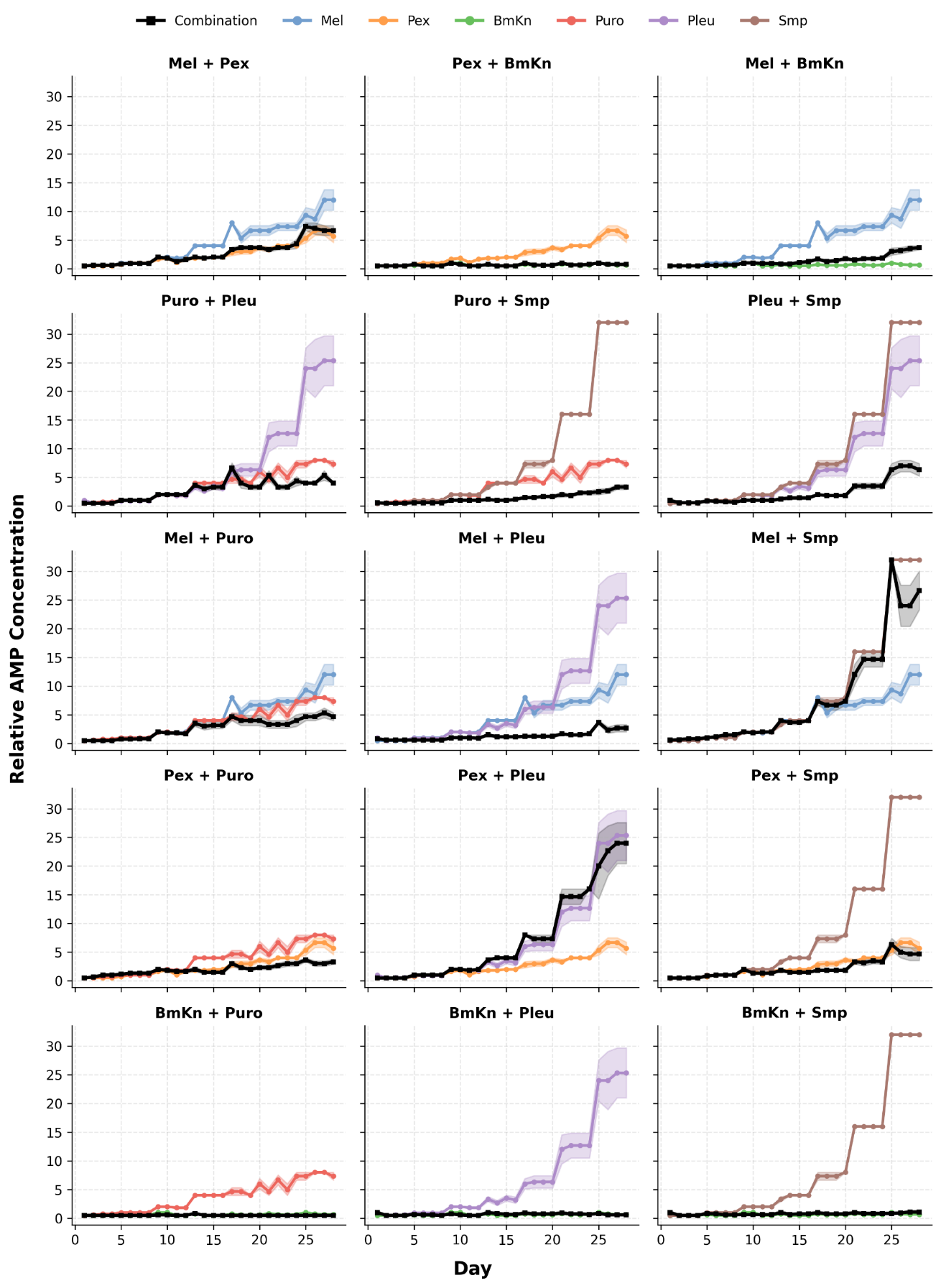

**Figure S6. Resistance evolution trajectories for all 15 pairwise AMP combinations.** Each panel shows the daily tolerated concentration (normalized to ancestral MIC) for one combination (black) alongside its two component single-AMP treatments (colored). Lines show means across six evolutionary replicates, with shaded areas representing standard error. Combinations are organized with membrane × membrane pairs (top row), intracellular pairs (second row), and mixed pairs (remaining rows).

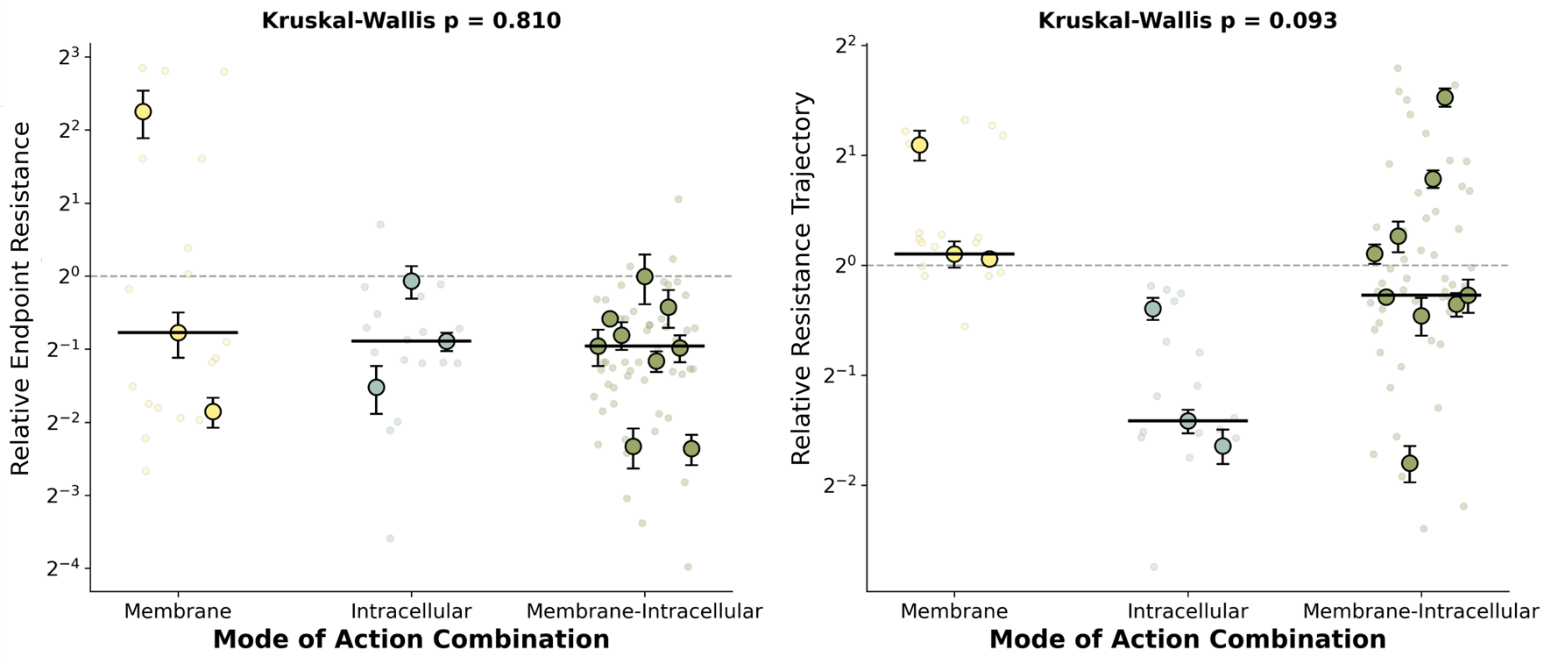

**Figure S7. Mode of action pairing does not predict combination efficacy.** Relative Endpoint Resistance (**left**) and Relative Resistance Trajectory (**right**) grouped by MOA pairing category: membrane × membrane (n = 3 combinations), intracellular × intracellular (n = 3), and membrane × intracellular (n = 9). Large colored points with error bars show combination-level means ± standard error. Small faded points show individual evolutionary replicates. Horizontal black lines indicate group medians. Dashed line at 1 indicates no difference from single-AMP controls. Neither metric differed significantly across MOA categories (Kruskal-Wallis p = 0.810 and p = 0.093, respectively).

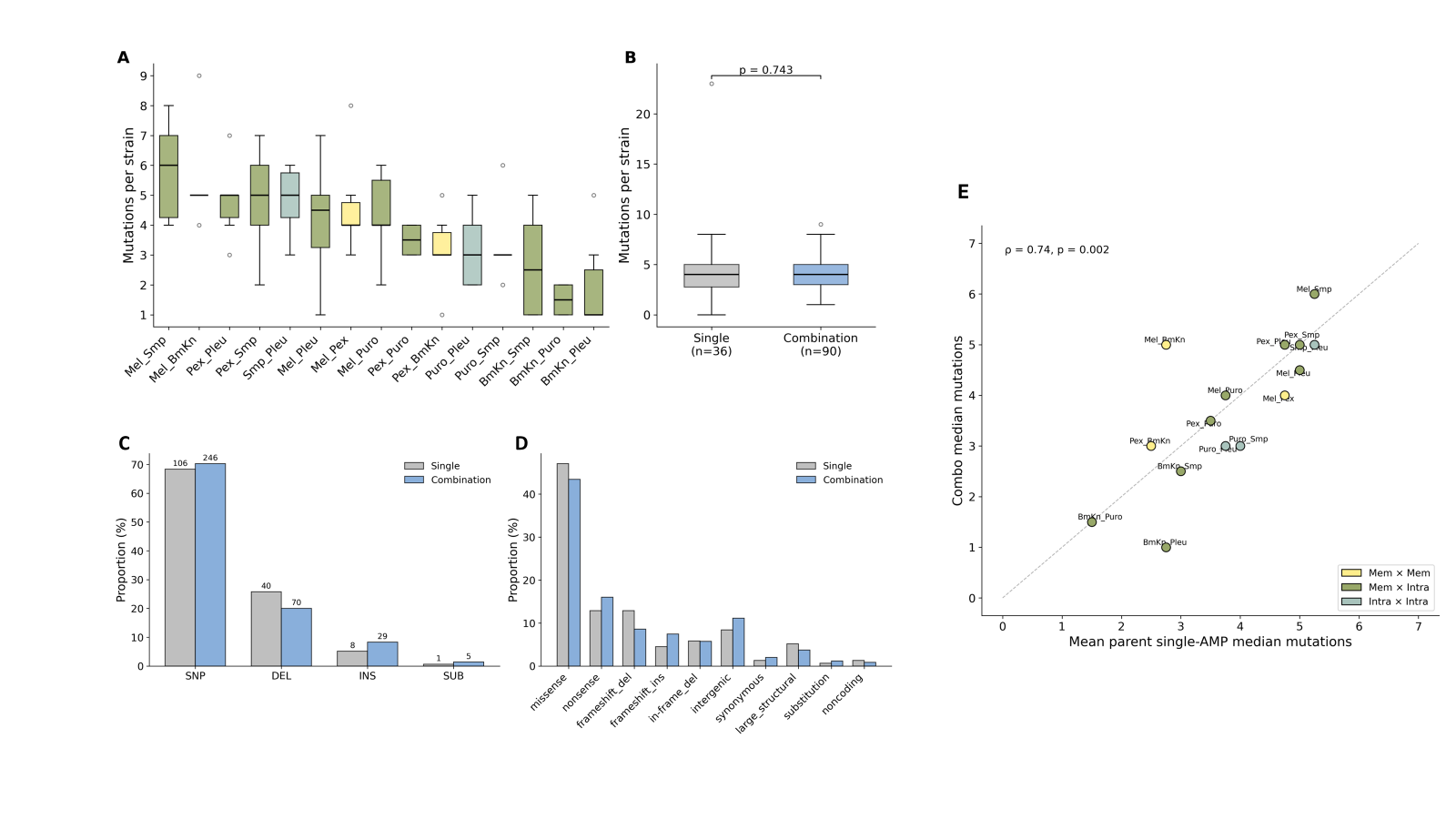

**Figure S8. Mutational landscape under combination therapy resembles single-AMP evolution in scale and character.** (**A**) Mutation burden per strain across all 15 combination treatments. Box colors indicate MOA pairing category (yellow, membrane pair; olive, membrane × intracellular; teal, intracellular pair). Combinations are sorted by median mutation burden. (**B**) Overall mutation burden comparison between single-AMP (n = 36) and combination-evolved (n = 90) strains. Median burden did not differ (Mann-Whitney p = 0.846). (**C**) Distribution of variant types in single-AMP (gray) and combination (blue) datasets. Numbers above bars indicate counts. The distributions were not significantly different (χ² = 3.40, p = 0.33). (**D**) Distribution of functional effects in single-AMP and combination datasets. The distributions were not significantly different (χ² = 2.92, p = 0.82). (**E**) Correlation between combination median mutation burden and the mean of the two parent single-AMP median mutation burdens. Each point represents one combination, colored by MOA pairing category. Dashed line shows y = x. Spearman ρ = 0.77, p = 0.001.

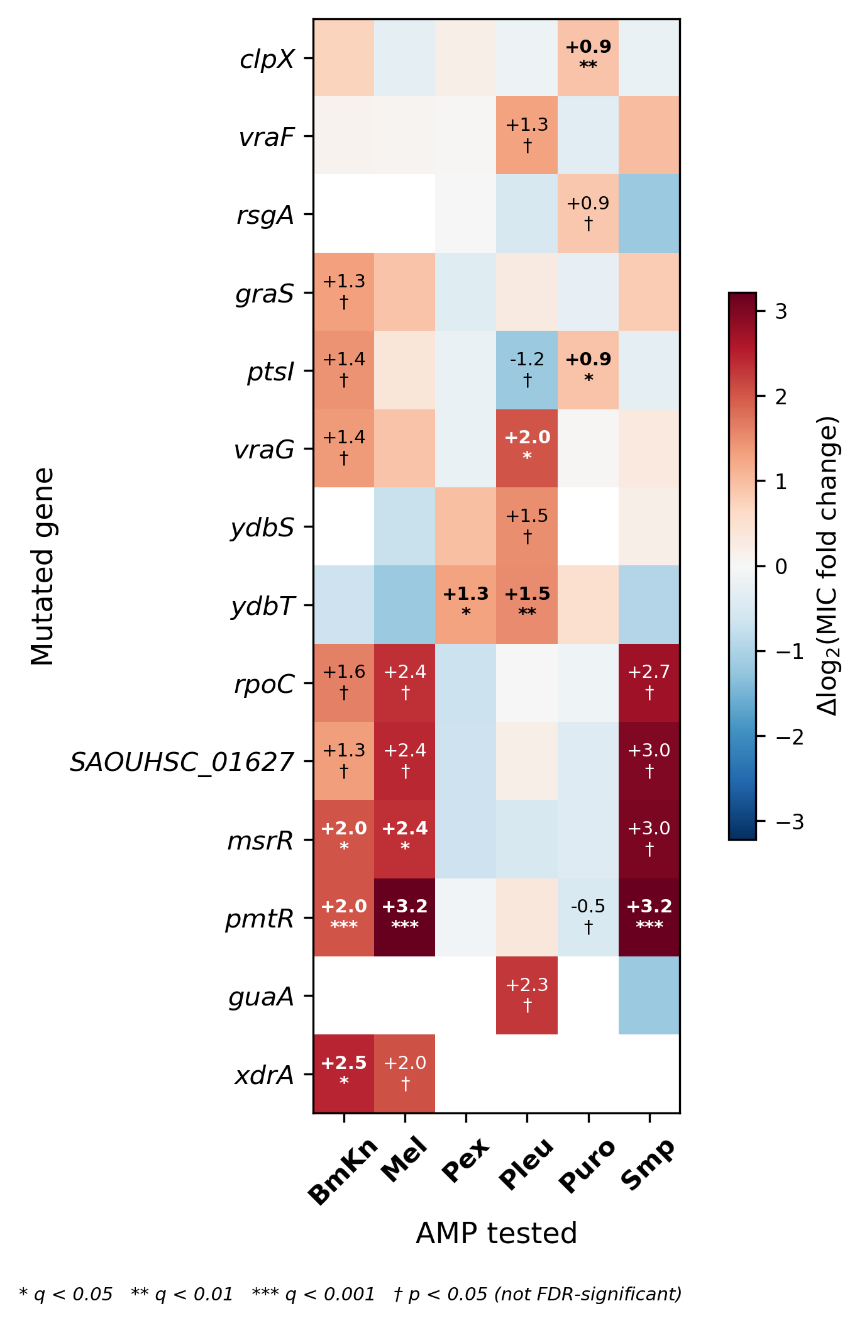

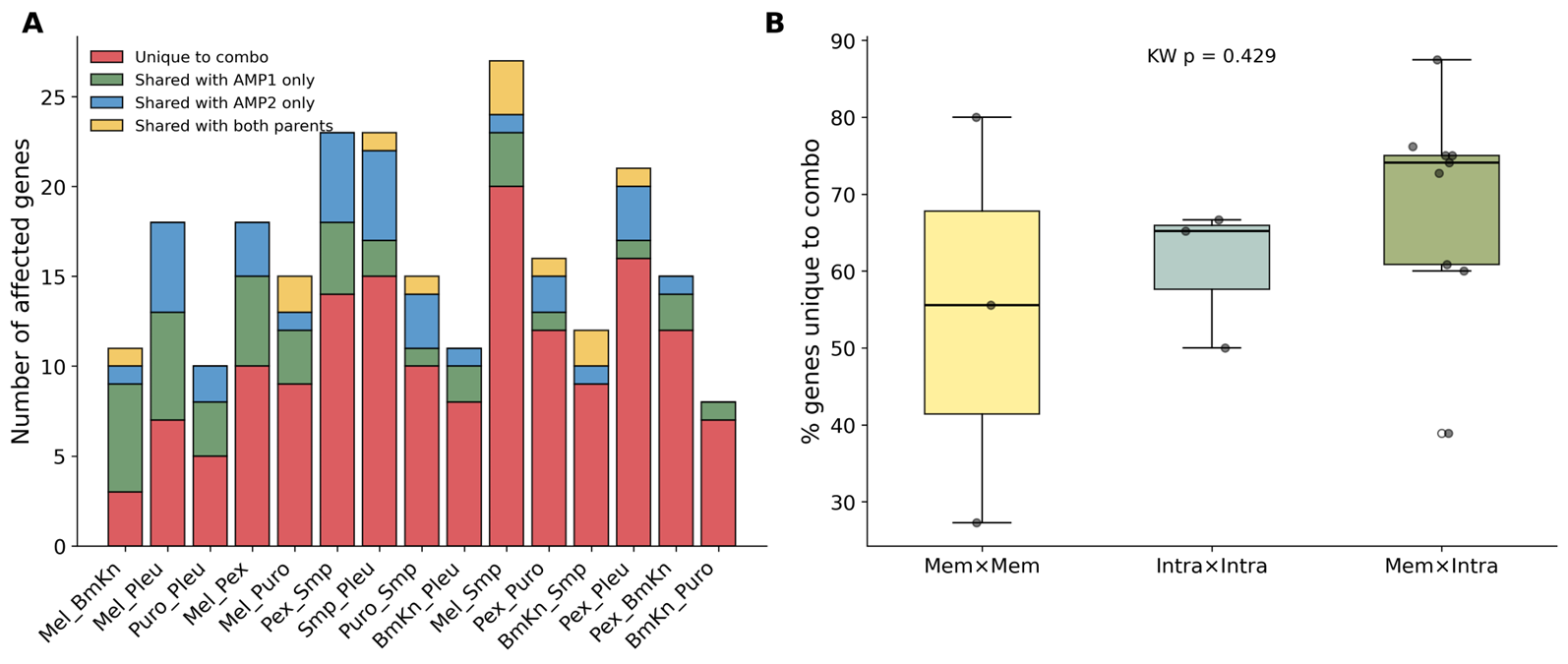

**Figure S10. Association between mutated genes and AMP-specific resistance.** Heatmap showing the mean difference in log₂ MIC fold-change between strains carrying a mutation in each gene (rows) and strains without that mutation, tested against each AMP (columns). Positive values (red) indicate that strains with the mutation show higher resistance to the tested AMP, and negative values (blue) indicate lower resistance. Annotations show effect sizes for significant associations only (* q < 0.05, ** q < 0.01, *** q < 0.001 after FDR correction, † p < 0.05 nominally significant but not FDR-corrected). Cells without annotations were not statistically significant.

**Figure S9. Gene-level novelty of combination-evolved mutational landscapes.** (A) Number of affected genes per combination treatment, categorized by whether they were unique to the combination (red), shared with only the first parent single-AMP lineage (green), shared with only the second parent (blue), or shared with both parents (yellow). Most affected genes in each combination were not mutated in either parent single-AMP lineage. (B) Percentage of affected genes unique to the combination, grouped by MOA pairing category. The proportion of unique genes did not differ across categories (Kruskal-Wallis p = 0.429).

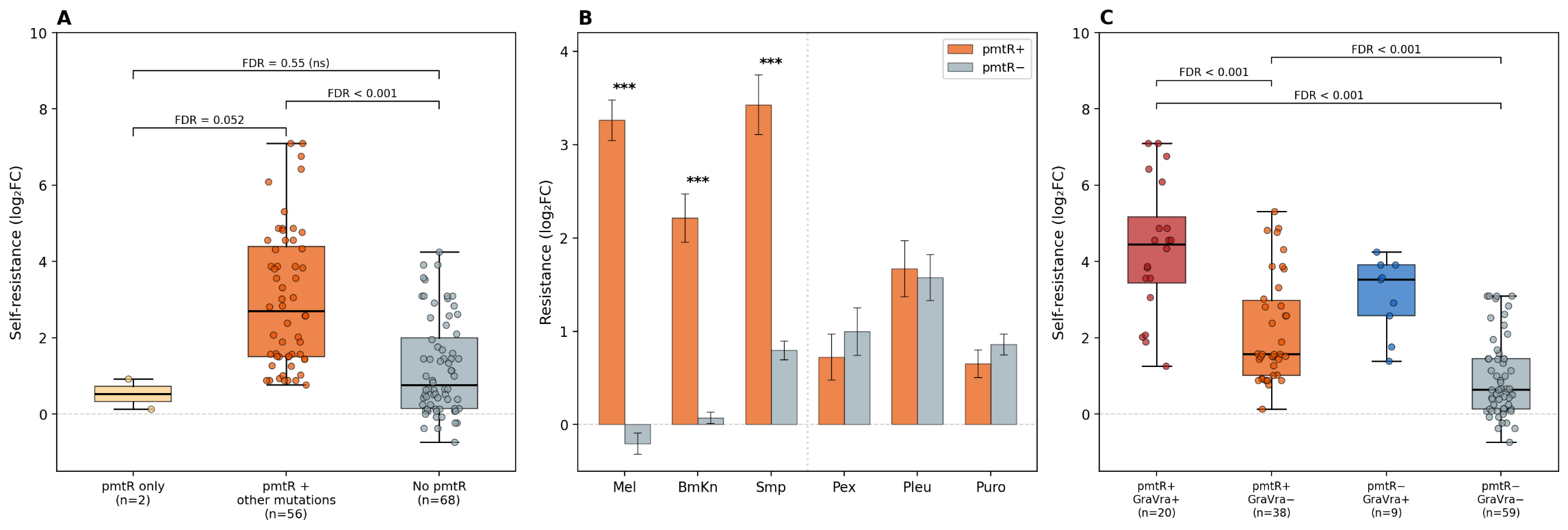

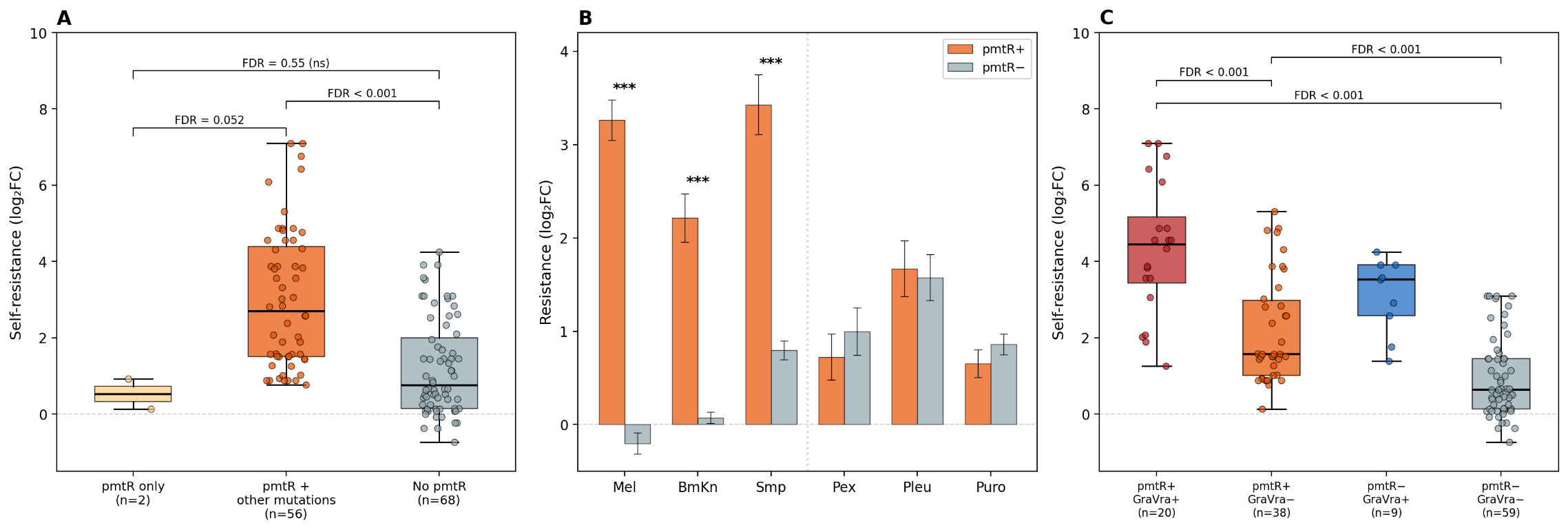

**Figure S11. pmtR inactivation primes resistance but requires additional mutations for high-level resistance.** (A) Self-resistance of all 126 evolved strains grouped by pmtR mutational status. pmtR-only strains (n = 2: BmKn5, BmKn_Puro1) carry no additional mutated genes. Self-resistance is defined as log₂ fold-change in MIC against the selecting AMP for single-AMP evolved strains, or as the mean log₂ fold-change against both component AMPs for combination-evolved strains. (B) Mean resistance of pmtR-positive and pmtR-negative strains tested against each AMP. Error bars show standard error of the mean (*** FDR < 0.001). The dotted line separates AMPs with significant pmtR association (Mel, BmKn, Smp) from those without (Pex, Pleu, Puro). (C) Self-resistance grouped by the presence or absence of pmtR and GraRS/VraFG network mutations (*vraF*, *vraG*, *graS*, *graR*, or *mprF*). The two mutation classes co-occur more frequently than expected by chance (Fisher's exact test, OR = 3.45, p = 0.006). Each dot represents one independently evolved lineage. Boxplots show median, interquartile range, and 1.5× IQR whiskers. All comparisons were performed using Mann-Whitney U tests with Benjamini-Hochberg FDR correction.

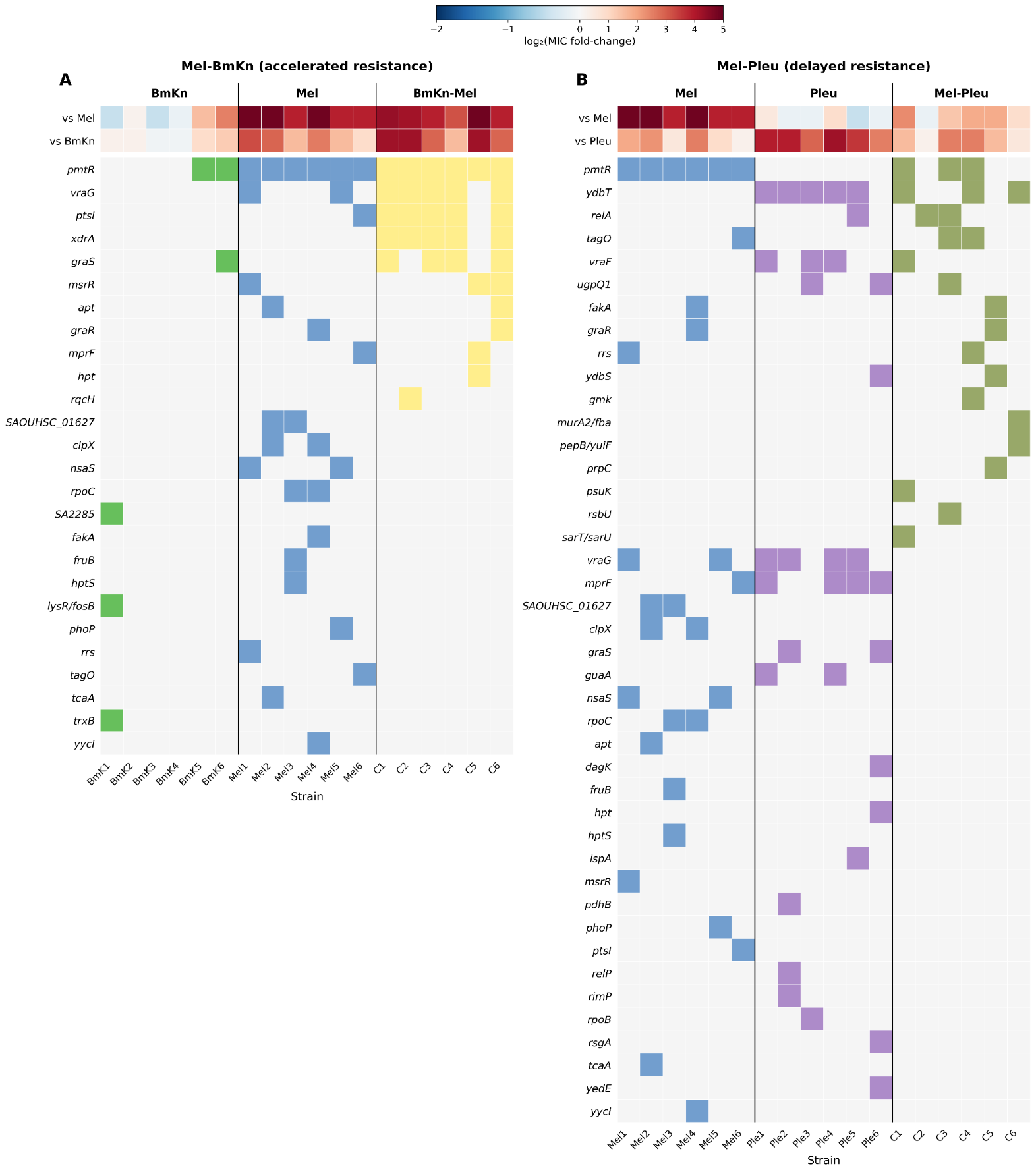

**Figure S12. Complete mutational profiles of Mel-BmKn and Mel-Pleu combination-evolved strains and their single-AMP counterparts.** Layout and color coding as in Fig. 6, with all mutated genes shown without frequency filtering. (A) Mel-BmKn and its component single-AMP treatments (BmKn, Mel). (B) Mel-Pleu and its component single-AMP treatments (Mel, Pleu).
